# Integrative analysis of the salt stress response in cyanobacteria

**DOI:** 10.1101/2021.07.28.454097

**Authors:** Stephan Klähn, Stefan Mikkat, Matthias Riediger, Jens Georg, Wolfgang R. Hess, Martin Hagemann

**Affiliations:** Helmholtz-Centre for Environmental Research - UFZ, Department of Solar Materials, Leipzig, Germany; University of Freiburg, Faculty of Biology, Genetics and Experimental Bioinformatics, Freiburg, Germany; Core Facility Proteome Analysis, Rostock University Medical Center, Rostock, Germany; University of Rostock, Institute of Biosciences, Dept. Plant Physiology, Rostock, Germany; Department Life, Light & Matter, University of Rostock, Rostock, Germany

**Keywords:** compatible solute, ion transport, regulatory RNAs, transcriptome-proteome correlation, salinity stress response

## Abstract

Microorganisms evolved specific acclimation strategies to thrive in environments of high or fluctuating salinities. Here, salt acclimation in the model cyanobacterium *Synechocystis* sp. PCC 6803 was analyzed by integrating transcriptomic, proteomic and metabolomic data. A dynamic reorganization of the transcriptome and proteome occurred during the first hours after salt shock, e.g. involving the upregulation of genes to activate compatible solute biochemistry balancing osmotic pressure. The massive accumulation of glucosylglycerol then had a measurable impact on the overall carbon and nitrogen metabolism. In addition, we observed the coordinated induction of putative regulatory RNAs and of several proteins known for their involvement in other stress responses. Overall, salt-induced changes in the proteome and transcriptome showed good correlations, especially among the stably up-regulated proteins and their transcripts. We define an extended salt stimulon comprising proteins directly or indirectly related to compatible solute metabolism, ion and water movements, and a distinct set of regulatory RNAs involved in post-transcriptional regulation. Our comprehensive data set provides the basis for engineering cyanobacterial salt tolerance and to further understand its regulation.

## Introduction

Salinity is a prominent environmental factor determining the natural distribution of microorganisms, as approximately 97% of the water resources contain more than 30 g salt (mainly sodium chloride) per liter. Accordingly, the capability of microorganisms to cope with high or changing salinities is crucial, not only in aquatic environments but also in terrestrial habitats, in which the alternation between evaporation and rainfall can rapidly change salt concentrations. In a hypersaline environment, in which the external salt concentration exceeds the cellular ion content, microorganisms have to manage two major challenges: (i) the low external water potential results in water loss from the cell and collapse of turgor pressure, and (ii) inorganic ions permeate into cells along the electrochemical gradient, which could compromise the structure of critical macromolecules. Accordingly, most microorganisms feature acclimation strategies aiming at maintaining a high water and low inorganic ion content in the cell. This so-called “salt-out” strategy is based on the active extrusion of inorganic ions accompanied by the accumulation of compatible solutes, i.e. highly soluble, non-toxic, low molecular-mass organic compounds, for osmotic equilibrium (e.g., Hagemann, 2011).

Cyanobacteria are a morphologically and physiologically diverse group of photoautotrophic bacteria that are found in nearly all light-exposed habitats including environments with different salinities such as freshwaters, oceans, and hypersaline ponds or soil surfaces in temperate and arid climates (Whitton and Potts, 2000). Previous studies of cyanobacterial salt tolerance revealed that those with low salt tolerance, mainly freshwater and soil cyanobacteria, accumulate the sugars sucrose and/or trehalose, those with moderate tolerance (mainly marine strains) synthesize the heteroside glucosylglycerol (GG), whereas halophilic strains (found in hypersaline habitats) usually contain glycine betaine (Reed et al., 1986; Hagemann, 2011). Some deviations from these preferences have been documented as well (e.g., Klähn et al., 2010a; Pade et al., 2012, 2016).

Cyanobacterial salt acclimation has been investigated in great detail using the model strain *Synechocystis* sp. PCC 6803 (*Synechocystis* 6803). This unicellular strain was originally isolated from a freshwater pond (Stanier et al., 1971). Nevertheless, *Synechocystis* 6803 represents a truly euryhaline organism able to grow in freshwater but also in media containing salt concentrations twice as high as in seawater (Reed and Stewart, 1985). The early availability of both, genetic tools (Grigorieva and Shestakov, 1982) and the complete genome sequence (Kaneko et al., 1996) established *Synechocystis* 6803 as photoautotrophic model organism. Using this strain, the molecular basis of salt-induced GG synthesis has been characterized, which is performed by the enzymes GG-phosphate synthase and GG-phosphate phosphatase, encoded by the genes *ggpS* (*sll1566*) and *ggpP* (*stpA*, *slr0746*), respectively (Hagemann et al., 1997; Marin et al., 1998). Today, *Synechocystis* 6803 is the best investigated photoautotrophic prokaryote represented by more than 3700 scientific publications, and has become popular as chassis for the introduction of pathways for the photosynthetic production of biofuels or chemical feedstock (e.g., Hagemann and Hess, 2018; Liu et al., 2019a). Salt-tolerant cyanobacterial strains such as *Synechocystis* 6803 also permit large scale cultivations in saline waters making the process more sustainable by avoiding competition for limited freshwater resources (Chisti, 2013).

Over the past two decades, omics technologies have been applied to study salt acclimation of cyanobacteria, particularly *Synechocystis* 6803. Genome-wide transcriptome analyses revealed the differential expression of hundreds of genes after sudden increases in the external salinity and allowed the identification of several potential regulatory proteins involved in their stress-induced expression (Kanesaki et al., 2002; Marin et al., 2003; Shoumskaya et al., 2005). However, most of these genes were only transiently induced or repressed. In long-term salt-acclimated cells, the expression of only 39 genes remained significantly enhanced (Marin et al., 2004). Later on, 2D-gel-based proteomics displayed a snapshot of the salt-regulated proteome, which identified 45 stably salt-induced soluble proteins (Fulda et al., 2006), while 20 proteins of the membrane fraction appeared to be salt-regulated (Huang et al., 2006). In the meantime, advanced omics technologies such as different RNA-seq technologies were established, which, for example, revealed that also large numbers of non-protein-coding RNAs (ncRNAs) are transcribed in *Synechocystis* 6803 (Mitschke et al., 2011; Kopf et al., 2014; Billis et al., 2014). Among them, two classes can be differentiated, cis-encoded antisense RNAs (asRNAs) transcribed from the opposite strand within protein-coding genes, and trans-encoded small regulatory RNAs (sRNAs) (Kopf and Hess, 2015; Georg and Hess, 2018). Some of the newly annotated asRNAs were demonstrated to act as regulators of their cognate mRNAs (Dühring et al., 2006; Sakurai et al., 2012) or sRNAs functioning in the acclimation response to changing environmental conditions (Georg et al., 2014, 2017; Klähn et al., 2015; Zhan et al., 2021). Moreover, tremendous progress has been made in the investigation of cyanobacteria using gel-free technologies for proteomics (Wegener et al., 2010; Gao et al., 2015a; Spät et al., 2021) or metabolomics (reviewed by Schwarz et al., 2013).

Here, we combined transcriptomic with proteomic and metabolomic approaches for a comprehensive characterization of the salt acclimation process in *Synechocystis* 6803. Previous studies (e.g., Marin et al., 2004; Fulda et al., 2006) showed that salt acclimation is a highly dynamic process, in which many genes/proteins showed an early but mostly transient response before long-term salt acclimation leads to stable, physiological meaningful changes in the gene/protein expression pattern. Therefore, we sampled *Synechocystis* 6803 cells at different time points up to 7 days after transfer from NaCl-free into medium containing 684 mM NaCl (equal to 4% NaCl). In addition to many salt-regulated proteins and their corresponding mRNAs, we identified several potentially regulatory asRNAs and sRNAs to be salt-stimulated as well. Finally, metabolomics revealed that the massive accumulation of the compatible solute GG has a broader impact on the overall primary carbon and nitrogen metabolism.

## Results

The response of *Synechocystis* 6803 to NaCl-induced hyperosmotic conditions (salt stress) was analyzed on transcriptome, proteome and metabolome levels at different time scales (Fig. 1). Changes in the expression profiles were first analyzed at a global scale. Then, selected examples were examined in a comprehensive way using all data sets.

**Figure 1.**
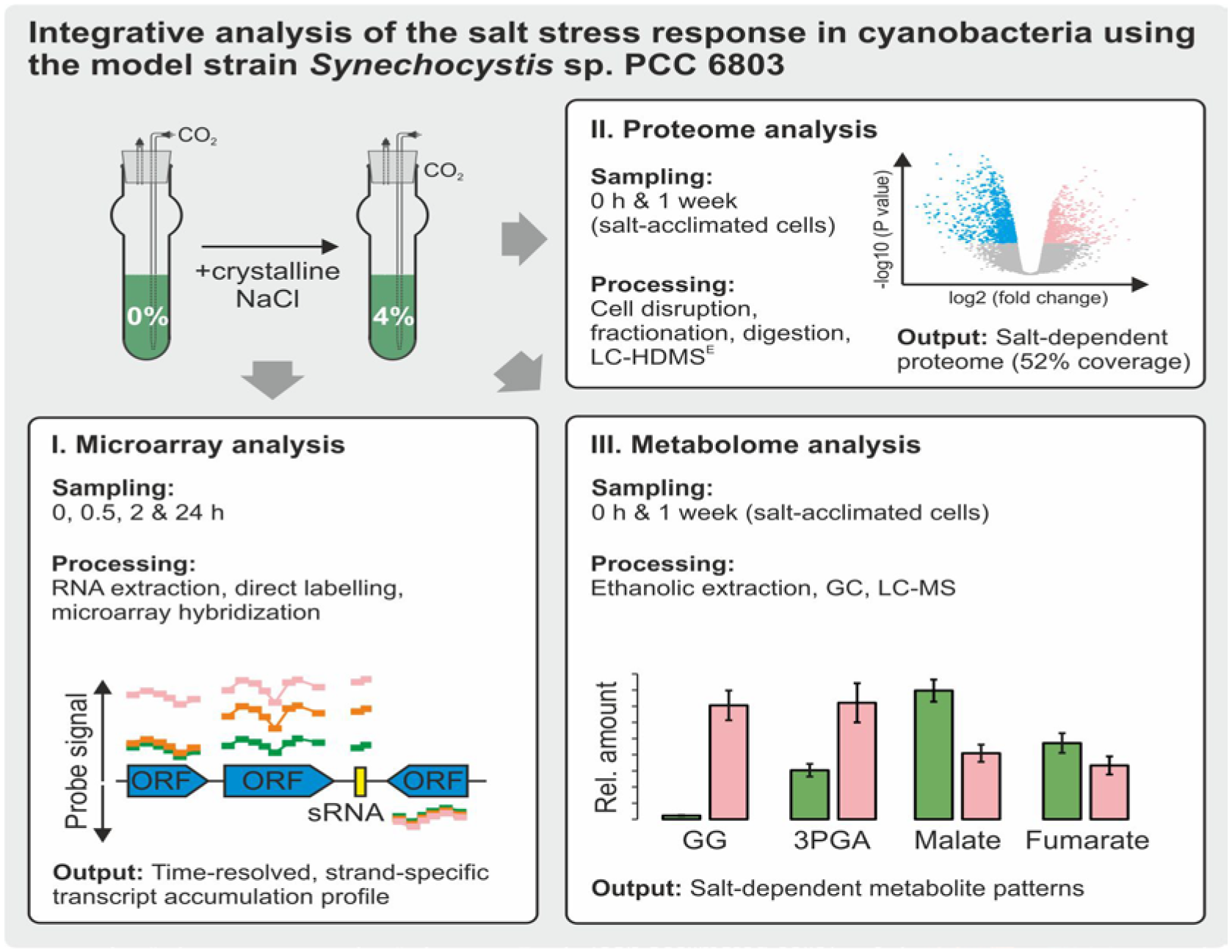
Overview on the applied approaches and conditions. Samples for microarray, proteome, and metabolome analyses were taken from cultures sparked with CO_2_-enriched air (5 %[v/v]). Sampling points for each experiment are given in the panels. Further details about cultivation, salt treatment and sample processing are given in the Materials & Methods section.

### Global transcriptome analysis

In the present study a microarray platform was used to detect RNA levels directly, without reverse transcription. This microarray contained probes for all protein-coding genes also covering so far non-annotated open reading frames as well as for ncRNAs such as cis-acting asRNAs and trans-acting sRNAs previously identified in *Synechocystis* 6803 (Mitschke et al., 2011; Kopf et al., 2014). In total the microarray covered 3364 mRNAs, 1940 asRNAs and 602 sRNA candidates. Previous studies analyzing the salt transcriptome of *Synechocystis* 6803 used DNA microarrays only covering 3079 mRNAs and no ncRNAs (Marin et al., 2004). Here, the microarray was hybridized with total RNA extracted from control cells (0% NaCl) and from cells exposed to 4% of NaCl for 0.5, 2, and 24 h. The time points were chosen to permit comparison with previously published microarray data (Marin et al., 2004). Gene expression changes along the *Synechocystis* 6803 chromosome are shown in the Suppl. Genome Plots.

For the selection of differentially expressed genes, we applied typical cut-off criteria, i.e. log_2_ fold change ≥│1│, p-value < 0.05. As the transcriptome composition was highly dynamic at the different time points, we first focused on protein coding genes (mRNAs). Compared to control conditions (i.e., cells grown in NaCl-free BG11 medium), several hundred mRNAs showed a changed abundance 0.5 and 2 h after salt addition, while after 24 h only 87 were up- and 31 down-regulated (Table 1; volcano plots of transcriptional changes are displayed in Suppl. Fig. S1), consistent with the previous report by Marin et al. (2004). Only a few mRNAs were significantly changed at all sampling points, i.e. 31 showed elevated levels and 19 were down-regulated.

**Table 1.** Global analysis of protein-coding genes showing altered transcript levels after salt shock of 684 mM NaCl. A gene was regarded as induced or repressed if the log_2_ fold change was higher or lower than 1 or −1 (*P* value < 0.05; n.a. – not analyzed).

|  | Time after salt shock | Marin et al., 2004 | Billis et al., 2014 | this study |
| --- | --- | --- | --- | --- |
| up-regulated mRNAs | 0.5 h | 652 | n.a. | 382 |
|  | 2 h | 477 | n.a. | 458 |
|  | 24 h | 48 | 133 | 87 |
| down-regulated mRNAs | 0.5 h | 329 | n.a. | 407 |
|  | 2 h | 268 | n.a. | 574 |
|  | 24 h | 48 | 368 | 31 |

To evaluate the microarray data set systematically, a cluster analysis was performed using *mfuzz* (Kumar and Futschik, 2007). Initially, 8538 transcript types were differentiated, including all mRNAs but also 5’UTRs, asRNAs, sRNAs and transcripts derived from start sites within genes; Suppl. Table S2). Two clusters (cluster 1 and 2) include transcripts that were induced and two other clusters (cluster 3 and 4) include transcripts that were repressed at specific time points after salt addition (Fig. 2A). Transcripts in cluster 1 and 3 peaked at 0.5 h, while transcripts in cluster 2 and 4 peaked at the 2 h time point. The majority of genes in all clusters, most pronounced in case of cluster 3, showed a clear tendency to return to the initial values at the end of the time course indicating that the short-term acclimation was complete.

**Figure 2.**
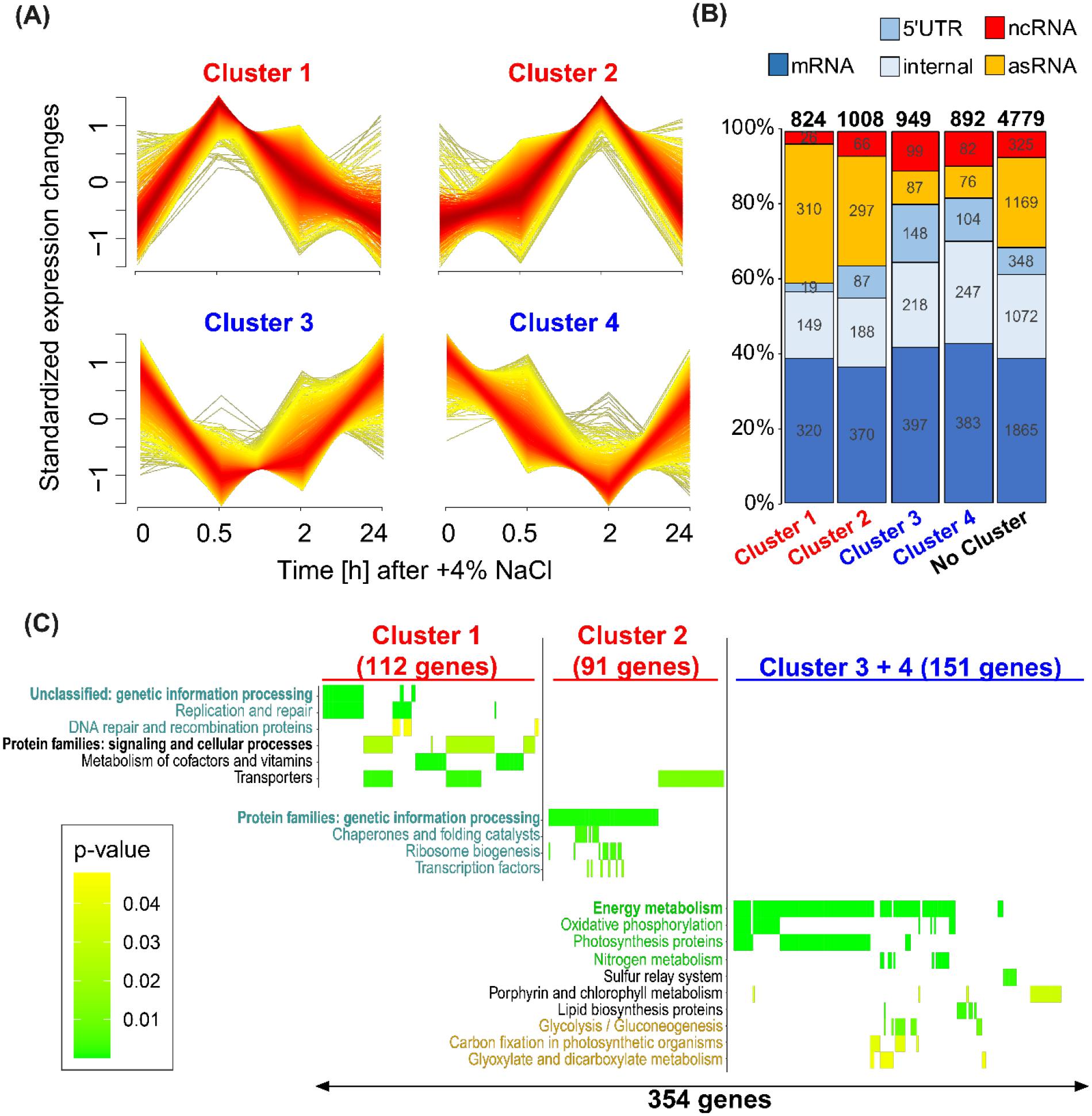
Cluster analysis of the salt stress microarray time series. **A.** Four major clusters of co-regulated transcripts were obtained. **B.** Distribution of different transcript types over the different clusters (see Suppl. Table S2 for the precise values and assignments). **C:** Functional enrichment analysis of proteins encoded by differentially regulated mRNAs according to KEGG Orthology (KO) terms for each cluster. Heatmap coloring represents the enrichment p-values (y axis = enriched KO terms, x axis = genes, see Suppl. Table S3 for detailed information).

**Table 2.**
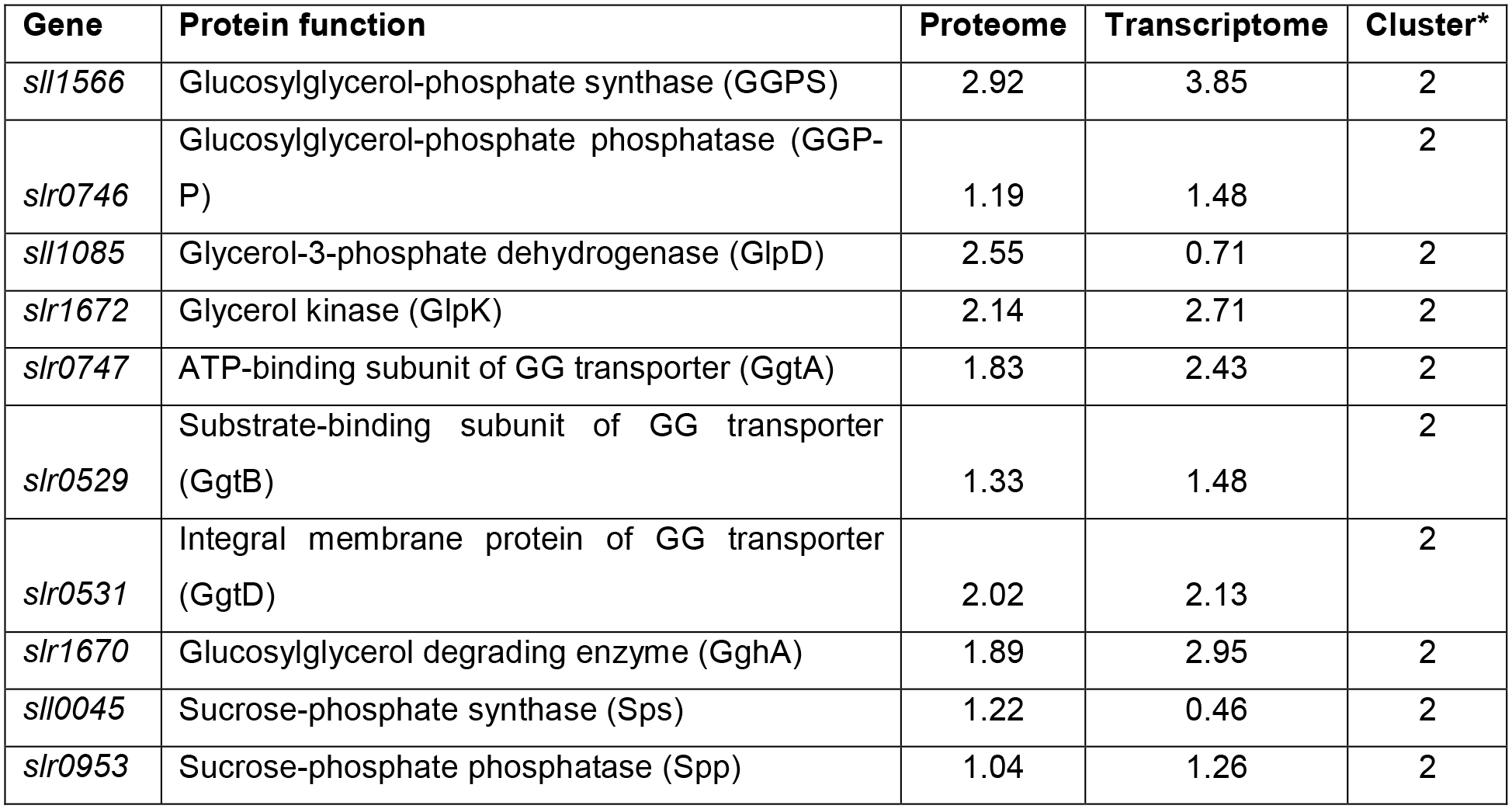
Expression of proteins involved in compatible solute metabolism and transport. (given are log_2_ fold changes of their levels in cells exposed for 7 d (proteome) or 24 h (transcriptome) to 684 mM NaCl versus control cells; *according to Cluster analysis shown in Fig. 2).

**Table 3.** Salt-regulated proteins that are involved in general stress tolerance. Given are log_2_ fold changes of their protein and corresponding mRNA levels in cells exposed for different times to 684 mM NaCl versus control cells.

| Gene | Protein function | Proteome | Transcriptome |  |  |
| --- | --- | --- | --- | --- | --- |
|  |  | 7 d | 24 h | 2 h | 0.5 h |
| *sl1863* | Unknown protein | 3.73 | 5.18 | 7.47 | 7.49 |
| *sl1862* | Unknown function | 3.30 | 4.00 | 5.28 | 5.30 |
| *sl11037* | Putative component of TRAP transporter | 3.01 | 0.63 | 0.17 | 0.38 |
| *slr0750* | Light-independent protochlorophyllide reductase subunit N | 3.00 | 1.29 | -1.88 | -0.86 |
| *slr0786* | Methionine aminopeptidase B | 2.83 | 4.77 | 5.32 | 0.95 |
| *slr0967* | Hypothetical protein, involved in stress responses | 2.43 | 3.00 | 3.28 | 0.27 |
| *Sl10528* | Putative zinc metalloprotease | 2.43 | 4.19 | 6.22 | 1.61 |
| *sl17064* | CRISPR-Cas system 2 | 2.18 | 0.57 | -2.56 | -1.00 |
| *sl10248* | Flavodoxin (IsiB) | 2.15 | 1.15 | 4.27 | 0.89 |
| *sl11988* | 33 kDa chaperonin (HSP33) | 2.06 | 0.42 | 0.21 | 0.35 |
| *sl10947* | Light-repressed protein A, LrpA | 1.83 | -1.69 | -2.54 | 0.84 |
| *slr2019* | Putative ATP binding subunit of ABC transporter | 1.79 | 2.13 | -0.33 | -0.11 |
| *slr1894* | General stress protein MrgA/Dps | 0.94 | 1.06 | 2.09 | 1.16 |
| *sl11514* | 16.6 kDa small heat shock protein, molecular chaperon | -1.68 | 0.91 | 5.85 | 4.25 |

Specific transcript types showed an uneven distribution in the different clusters, i.e. clusters 1 and 2 contain a significantly higher number of asRNAs and lower number of sRNAs while cluster 3 and 4 showed the opposite. Protein-coding transcripts are more present in the downregulated clusters (780 mRNAs) than in the upregulated clusters (690 mRNAs) (Fig. 2B). Functional enrichment analysis of the proteins encoded by mRNAs in each cluster was performed according to annotations from KEGG Orthology (KO) terms. On the one hand, mRNAs within the rapidly induced cluster 1 are enriched in proteins associated to replication and repair, cofactor biosynthesis, signaling, and transport, while the mRNAs of cluster 2 are enriched in protein families associated to transport, genetic information processing such as chaperones and folding catalysts, ribosome biogenesis, or transcription factors (Fig. 2C). On the other hand, transiently repressed mRNAs in the clusters 3 and 4 showed similar functionalities, and thus were analyzed jointly. The most pronounced functional enrichment could be seen for transcripts encoding proteins associated to energy and metabolism, such as oxidative phosphorylation, photosynthesis, nitrogen metabolism, or related metabolic functions, such as pathways of porphyrin and chlorophyll metabolism, lipid biosynthesis or pathways for carbon metabolism.

#### Salt-regulated asRNA:mRNA pairs

Most of the salt-induced changes of asRNA levels were transient, consistent with the observations for mRNAs. Our analysis considered only asRNAs that overlap on the opposite strand with the respective mRNAs, which led to the identification of 79 inversely regulated and 82 co-regulated asRNA/mRNA pairs (Fig. 3) at a Pearson correlation coefficient ≥│0.65│ (details in Table S4). Previous work in *Synechocystis* 6803 showed that both modes of regulation can be functionally relevant (Dühring et al., 2006; Eisenhut et al., 2012; Sakurai et al., 2012).

**Figure 3.**
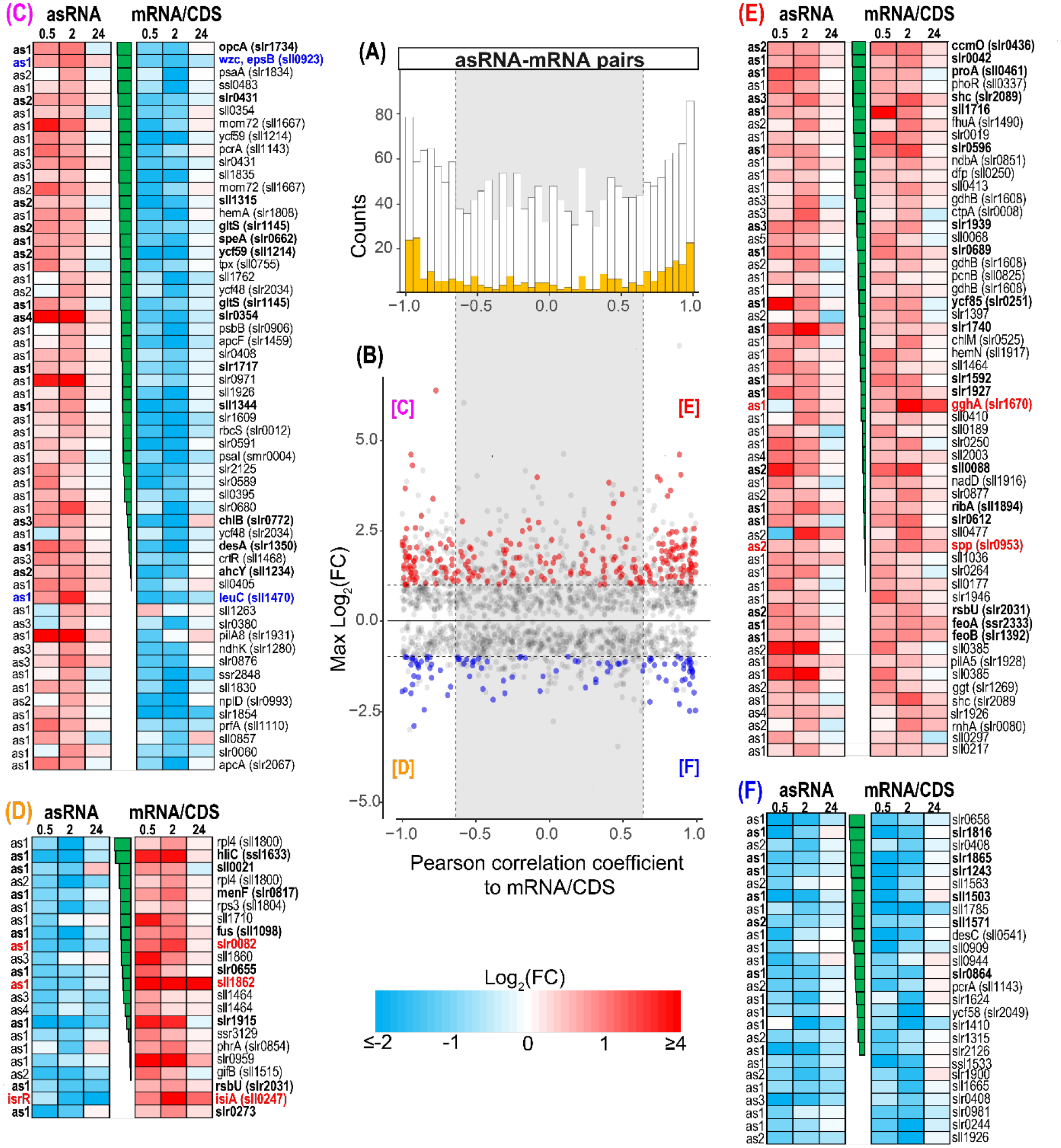
Comparison of salt-regulated mRNA:asRNA pairs with inverse or similar induction patterns. **A.** Histogram of Pearson correlation coefficients for expression profiles of all detected asRNA/mRNA pairs. Highlighted are asRNA/mRNA pairs, which both were assigned to an expression cluster with significant differential expression at least at one-time point. **B.** Scatterplot of maximum absolute log_2_ fold changes for every asRNA vs. Pearson correlation coefficients to its cognate mRNA. Colored points show the expression cluster assignment for the respective asRNA (red = cluster 1 + 2; blue = cluster 3 + 4). White backgrounds indicate strong correlation between asRNA/mRNA pairs (either ≥0.65 or ≤-0.65). Numbers of individual cluster assignments for asRNA/mRNA pairs are indicated in the boxes. **C.** 57 pairs were asRNA-induced and mRNA-repressed, **D.** 22 pairs were asRNA-repressed and mRNA-induced, **E.** 56 pairs were asRNA- and mRNA-induced, **F.** 26 pairs were asRNA- and mRNA-repressed. Only asRNA/mRNA pairs are given, which both were assigned to an expression cluster with a Pearson correlation coefficient of either ≥0.65 or ≤-0.65 at least at one-time point. The heat maps illustrate the log_2_(FC) at the individual measurements from the microarray experiments for asRNAs (left) and mRNAs (right). The heatmaps are sorted according to Pearson correlation coefficient (center). Examples that are mentioned in the text are highlighted based on their expression. Bold font indicates at least two significant differential expression measurements per transcript. Details of the asRNA/mRNA analysis are given in Suppl. Table S4.

**Table 4.** Salt effects on proteins involved in central carbon metabolism. (given are log_2_ fold changes of their levels in cells exposed for 7 d (proteome) or 24 h (transcriptome) to 684 mM NaCl versus control cells)

| Gene | Protein function | Proteome | Transcriptome |
| --- | --- | --- | --- |
| *slr0394* | Phosphoglycerate kinase | 0.76 | 0.58 |
| *slr0884* | Glyceraldehyde-3-phosphate dehydrogenase 1, Gap1 | 0.75 | 1.35 |
| *sl10842* | Neopullulanase, NpIT | 1.28 | 0.83 |
| *slr0237* | Glycogen debranching enzyme 1, GlgX1 | 1.16 | 1.13 |
| *slr1857* | Glycogen debranching enzyme 2, GlgX2 | -1.12 | -1.12 |
| *slr1367* | Alpha-1,4 glucan phosphorylase | 0.61 | 0.81 |
| *sl1356* | Glycogen phosphorylase 2, GlgP2 | 0.08 | 0.53 |
| *sl10587* | Pyruvate kinase 1 (PK 1) | 0.76 | 0.60 |
| *sl1275* | Pyruvate kinase 2 (PK 2) | -0.45 | -0.78 |
| *slr0301* | Phosphoenolpyruvate synthase | -1.19 | -0.97 |

Among the asRNA/mRNA pairs is *sll1862* (Fig. 3D) encoding one of the most abundant salt shock proteins (see below). The massive accumulation of stress protein Sll1862 is correlated by the inversely related levels of its mRNA versus asRNA. Another asRNA seems to be involved in controlling the salt-stimulated expression of *slr0082* (Fig. 3D), which encodes the ribosomal protein S12 methylthiotransferase RimO. This gene was early detected to be salt-induced by subtractive hybridization (Vinnemeier and Hagemann, 1999). It is transcribed in an operon together with the RNA helicase CrhR, which has been shown to be involved in multiple stress responses and is regulated by redox changes (Ritter et al., 2020). Moreover, the mRNA level of *sll0923* encoding the tyrosine kinase Wzc was less abundant after salt addition (Wzc protein level was lowered with a FC of 0.53), while the corresponding asRNA showed increased abundance (Fig. 3C). Wzc is involved in the synthesis of extracellular polysaccharides (EPS; Pereira et al., 2019), hence, the lowered expression of *sll0923* is consistent with observation of reduced EPS synthesis in salt-acclimated *Synechocystis* 6803 cells (Kirsch et al., 2017). Among the 56 mRNA/asRNA pairs showing similar induction patterns, the gene *slr0953* encoding sucrose-phosphate phosphatase showed higher mRNA as well as asRNA levels (Fig. 3E). It could be assumed that in this case the asRNA supports the stability of the *slr0953* mRNA contributing to the transiently elevated sucrose levels in salt-treated cells.

Moreover, we identified also not previously known salt-regulated transcripts, e.g. the *sll1470* asRNA (Fig. 3). Although this asRNA originates from a TSS downstream of *sll1470*, it overlaps the 3’ end of the gene and a clear anti-correlation with the transcript accumulation of *sll1470* during salt shock was detected (Fig. 3C). Gene *sll1470* encodes the large subunit of 3-isopropylmalate dehydratase, an important enzyme connecting pyruvate metabolism with leucine/isoleucine/valine biosynthesis. Therefore, its lowered expression during early salt shock likely contributes to the metabolic reorganization towards synthesizing GG consistent with the lowered valine accumulation (see below).

#### Salt-regulated sRNAs

The used microarray also permitted to search for salt-induced changes in the abundance of sRNAs, which are in contrast to asRNAs encoded in trans. Interestingly, according to the cluster analysis (Fig. 2), both ncRNAs types responded differently to the salinity increase: asRNAs were mainly upregulated (cluster 1 and 2) while sRNAs were mainly downregulated (cluster 3 and 4). The salt-induced sRNA patterns, were again highly dynamic and mainly transient. For example, the sRNA IsaR1 was found to be transiently up-regulated in response to salt but returned to control levels in long-term salt acclimated cells (Suppl. Table S1). IsaR1 is a key player in the acclimation of the photosynthetic apparatus to iron starvation by targeting several mRNAs encoding Fe^2+^-containing proteins and enzymes involved in pigment and FeS cluster biosynthesis (Georg et al., 2017). In addition to its role in iron acclimation, IsaR1 controls directly the synthesis of a key enzyme in GG synthesis, GgpS at the post-transcriptional level early during salt acclimation (Rübsam et al., 2018).

Altogether, eleven sRNA candidates were found to be differentially expressed at all-time points, three candidates were up- and 8 were down-regulated (Suppl. Table S5). Their potential roles in the salt acclimation process is an interesting topic for future research.

#### Global evaluation of the proteome in salt-acclimated cells

Proteome analyses using data-independent acquisition mass spectrometry (HDMS^E^) were performed with extracts from cells cultivated at either 0% NaCl (control cultures) or 4% NaCl (7 days’ salt-acclimated cultures, Fig. 1). To improve coverage and to obtain additional information on cellular localization, the proteome of different fractions was analyzed: 1. Total protein extracts obtained without any centrifugation; 2. The debris fraction obtained after low speed centrifugation (yellowish brown pellet); 3. The soluble fraction representing the blueish supernatant after high-speed centrifugation; and, 4. The membrane-enriched fraction representing the washed green pellets after high-speed centrifugation (further details are described in the supplementary material).

In total, 1816 proteins, approximately 52% of the entire proteome of *Synechocystis* 6803 in the UniProt database, were identified by at least two tryptic peptides per protein (Suppl. Table S6). In particular, 1253 proteins were found in the total extracts and 823 in the soluble fraction, while 1608 and 1421 proteins were identified in the membrane-enriched and debris fractions, respectively (Fig. 4). Similar to previous reports on the *Synechocystis* 6803 proteome (Gao et al., 2015a), many proteins (79%) were found in both, the soluble as well as the membrane fractions, although the membrane pellet was washed with high salt and high pH buffers. The occurrence of proteins in different fractions clearly correlates with their numbers of transmembrane helices (Suppl. Fig. S2). In the present study, we also investigated the cell debris that is usually removed from proteome analyses if cellular fractionation is performed. The vast majority (96.3%) of proteins in this fraction was also identified in the membrane-enriched fraction. However, 29 proteins were exclusively found in the debris fraction, for example the large Slr1567 protein, a putative outer membrane protein (Suppl. Table S7). Many other proteins enriched in the debris fraction are annotated as components of the cell envelope or of the outer membrane-bound periplasmic space.

**Figure 4.**
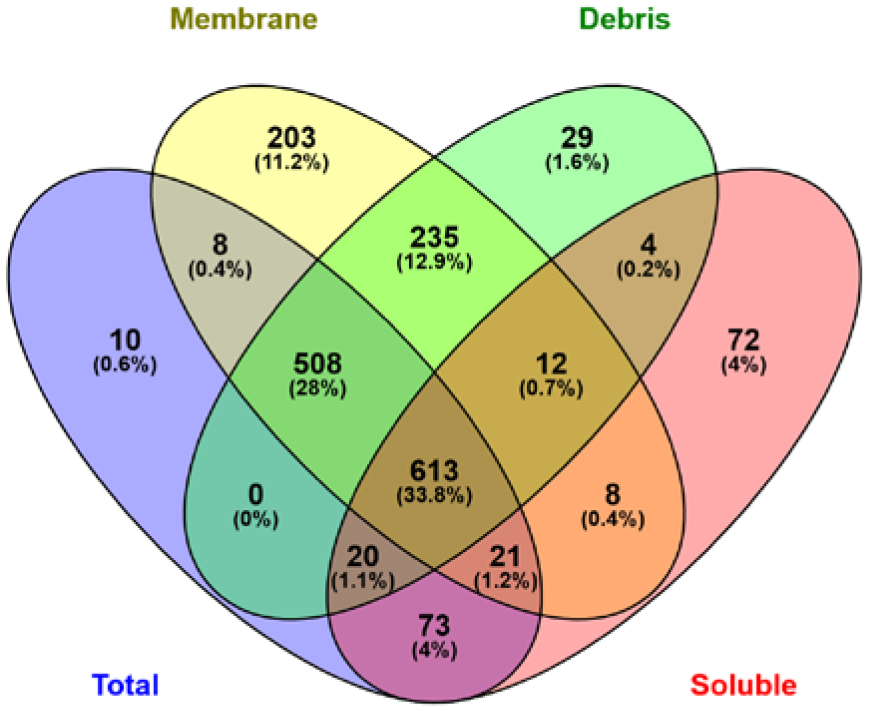
Overlap of the proteome among the different protein fractions. Venn diagram showing numbers and percentages of identified proteins in the total protein extract and in the subcellular fractions of debris, soluble or membrane proteins.

Salt-dependent changes in protein abundances were evaluated in total protein extracts as well as three subcellular fractions (Fig. 4). For the selection of differentially expressed proteins cut-off criteria were defined by a fold change of ≥│1.5│ and a corresponding p-value < 0.05. However, for a large number of proteins less pronounced fold change values were also statistically significant. The volcano plots indicate that the group of stably up-regulated proteins (Suppl. Fig. S3) displayed larger fold changes than down-regulated proteins, resembling the observations made with the transcriptomic data (Suppl. Fig. S1).

Next, we analyzed whether or not particular proteins showed consistent salt-related changes in the different proteome fractions. For most proteins, well-matching values were found in the different fractions and in the total extract. However, in some cases the relative abundances showed an inverse relation in membrane and soluble fractions. For example, many ribosomal proteins occurred in lower amounts in the soluble fraction but were elevated in the membrane and debris fractions of salt-acclimated cells compared to control cells (Suppl. Fig. S4). Similar patterns were found, among others, for the subunits of the ATP synthase, the phycobilisome linker polypeptides, and the RNA polymerase. These results indicate cultivation-dependent differences in the distribution of proteins to the different cellular fractions as reported in other studies (Gao et al., 2015b; Pattanayak et al., 2020). In our case, some protein complexes such as ribosomes might be more stable under high salt conditions, resulting in their enrichment in the membrane fraction and depletion in the soluble fraction. To deal with this situation, we calculated a weighted fold change from the subcellular fractions data of each protein and showed that it was highly concordant with the corresponding fold change from the total protein extract (Suppl. Fig. S5; more detailed 8 description is given in the supplementary material). Finally, the mean value of the weighted fold change from the subcellular fractions and the fold change from the total protein extract was used. If only one of these two values was available, it was used directly as final fold change. This led to a list of 1803 proteins, to which two membrane proteins were appended that are known (Slr0531) or suspected (Sll1037) to be important for salt acclimation, but were identified by individual peptides only. As a final result, 190 proteins were up-regulated and 189 protein down-regulated 7 days after salt shock among the 1805 quantified proteins (Suppl. Table S6).

#### Correlation analysis of salt-stimulated transcriptome and proteome

A correlation analysis was performed to analyze the overall relation between transcriptomic (log_2_ fold changes of mRNA after 24 h) and proteomic changes (log_2_ fold changes of protein abundances after 7 d). 1749 transcript/protein pairs could be matched (Fig. 5; Suppl. Table S8). The Pearson correlation coefficient for the proteomic and transcriptomic data sets was r = 0.58 indicating a quite good relationship, especially taking into consideration that sampling was done in different laboratories and at different time points. To grade the correlation between the newly acquired transcriptomic and proteomic data sets, we also compared our proteome data to the previously published transcriptome data sets by Marin et al. (2004) and Billis et al. (2014). In both cases, lower correlation coefficients of r = 0.41 and r = 0.42, respectively, were obtained. Next, we calculated the correlation coefficients between our transcriptome data and the transcriptome data of Marin et al. (2004) and Billis et al. (2014) using the same subset of mRNAs as for the comparison with the proteome. Surprisingly, the obtained coefficients of r = 0.49 and r = 0.50 showed a slightly lower correlation between different transcriptomic data sets than correlation between the proteomic and the present transcriptomic data set (Fig. 5C). These findings indicate that culture conditions significantly influence the comparability of different data. Nevertheless, in all cases a close correlation was observed for the expression of genes that are of direct importance for salt acclimation, while the expression of other genes can vary depending on small differences between the culture conditions leading to relatively low Pearson correlation coefficients.

**Figure 5.**
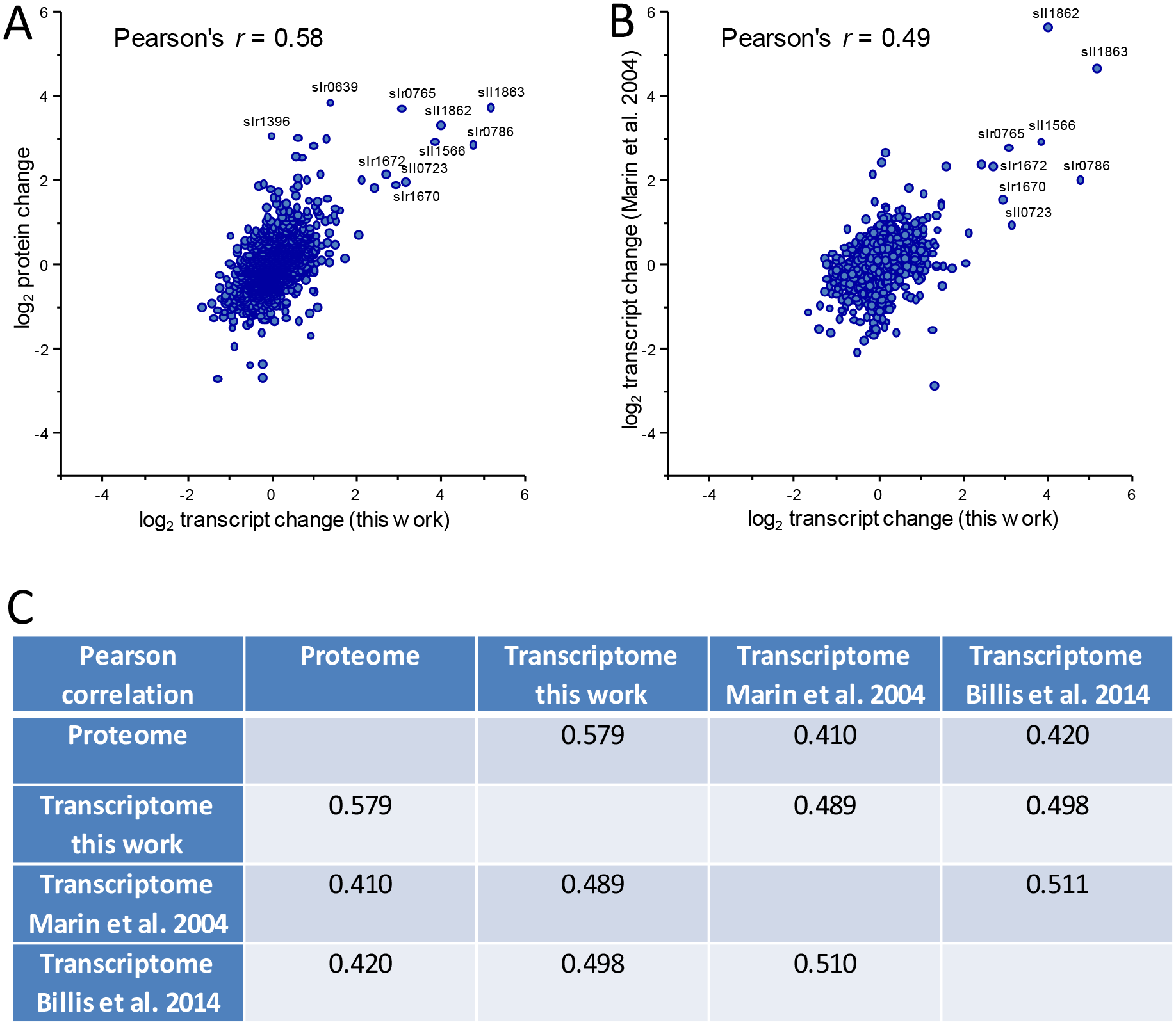
Correlation between transcriptome (24 h) and proteome (7 d) dynamics in response to high salt conditions. **A.** Data originate from the present study. **B.** The newly obtained proteome data were compared with the previous transcriptomic study (Marin et al., 2004). Scatterplots display the correlation of protein and transcript ratios. 1749 transcripts with reported fold changes could be mapped to corresponding changes in protein abundances. Matches with ratio differences below log_2_ 1.5-fold changes were considered to be similar. **C.** Table displaying Person’s correlation coefficients between the different data sets.

### Detailed examination of specific processes

#### Compatible solute metabolism and transport constitutes a salt-specific stimulon

Central to salt acclimation of *Synechocystis* 6803 is the accumulation of the main compatible solute GG, because mutants affected in the genes *ggpS* and *ggpP* (*stpA*) encoding the GG synthesis enzymes showed the highest degree of salt sensitivity compared to wild type (Hagemann et al., 1997; Marin et al., 1998). As initially found by Reed and Stewart (1985), salt-acclimated cells accumulate high amounts of the compatible solute GG, which represents the by far largest pool of low molecular mass organic compounds. The amount of GG is approximately 2000times higher in salt-grown cells compared to the trace amounts of GG in control cells (Fig. 6C). The second compatible solute sucrose is approximately 1000-fold less abundant (0.28 nmol OD_750_^-1^ ml^-1^) than GG in salt-acclimated cells (Suppl. Table S10), because it mainly plays a role as transiently accumulated osmolyte after salt shocks in *Synechocystis* 6803 (Kirsch et al., 2019).

**Figure 6.**
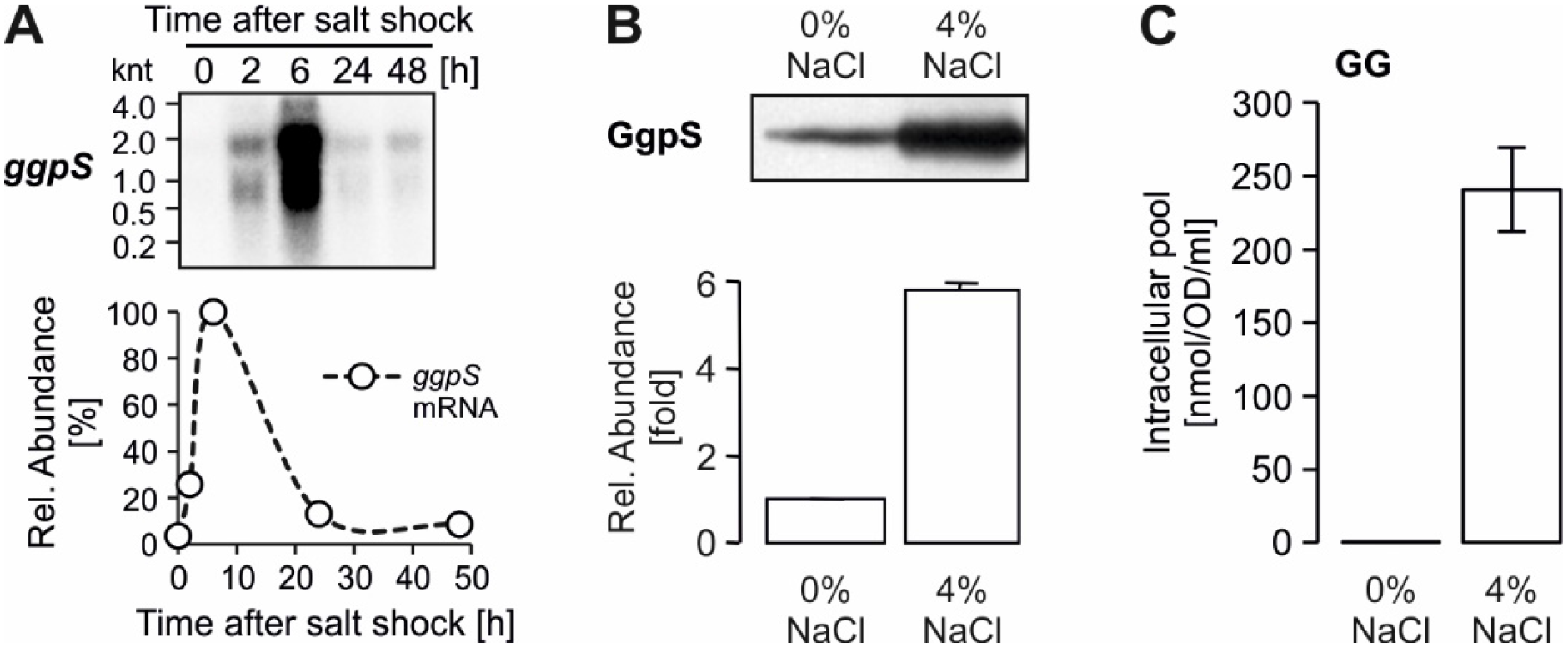
Salt-dependent up-regulation of the GG synthesis key enzyme, GG-phosphate synthase (GgpS). **A.** Northern-blot showing accumulation kinetics of the *ggpS* transcript in response to salt shock of 4 % [w/v] NaCl. Relative abundances were calculated after densitometric evaluation of the blot signals (signal obtained for 6 h was set as maximum, 100%). **B.** Western blot confirming increased GgpS abundance in cells acclimated to 4% NaCl. Relative abundance was calculated after densitometric evaluation of blot signals (signal from control cells was set as 1). Data are the mean ± SD of values obtained from three individual blots. **C.** Intracellular accumulation of GG in cells acclimated for 7 days to 4% NaCl. Data are the mean ± SD of 6 replicates.

Corresponding to the high GG accumulation, the GgpS and GgpP proteins and mRNAs showed significantly elevated levels (Table 2), which is supported for GgpS by Northern- and Western-blotting (Fig. 6AB). The sucrose synthesis enzymes Sps and Spp also exceeded the threshold for significant protein changes (Table 2), whereas the *sps* mRNA was only transiently increased (Suppl. Table S1) consistent with the transient accumulation profile of sucrose (Kirsch et al., 2019). The *ggpS* downstream gene *glpD* encoding glycerol 3-phosphate dehydrogenase, which is involved in the synthesis of the GG precursor glycerol 3-phosphate (G3P) from dihydroxyacetone phosphate, is also salt-stimulated. Overlapping with the *ggpS* promoter region exist the small ORF *ssl3076* that encodes for the *ggpS* repressor GgpR (Klähn et al., 2010b), which could not be detected in the proteome. Upstream of *ggpS* on the opposite strand, the genes for the GG hydrolase (GghA) and glycerol kinase (GlpK), the latter is involved in synthesis of the GG precursor G3P from glycerol, are located. They show similar expression pattern as *ggpS*. In contrast to the salt induction of genes for G3P synthesis, the *glgC* gene encoding the enzyme for ADP-glucose synthesis, the second precursor for GG, is not salt-regulated on RNA or protein levels.

Immediately downstream of *ggpP*, the salt-induced GgtA protein is encoded that acts as the ATP-binding subunit of the GG transporter (Ggt). The genes for the other Ggt subunits, GgtB, C and D (two of them were identified among the salt-stimulated proteins) form a separate salt-induced operon. The co-regulation of genes and proteins for GG synthesis and the ABC-type osmolyte transporter Ggt, which all belong to the cluster 2 (Table 2), indicates their functional cooperation in salt-acclimated cells, in which the transporter is mainly responsible for the avoidance of GG (and sucrose) leakage from the cells (Mikkat and Hagemann, 2000). However, the GG hydrolase GghA (Slr1670), which degrades GG into glucose and glycerol after hypo-osmotic treatments (Kirsch et al., 2017), showed also enhanced expression on mRNA and protein level in salt-acclimated cells. It has been discussed that the GG synthesis and degradation are mainly regulated at enzyme activity level that are differentially affected by cellular ion contents; elevated internal ion content leads to biochemical activation of GG synthesis and inactivation of GG cleavage (Kirsch et al., 2019). Hence, the increased amounts of putatively biochemically inactive GghA protein likely prepares *Synechocystis* 6803 cells for sudden decreases in osmolarity. It should be noted that the GgpP and GlpD proteins involved in GG synthesis were not detected in the soluble but exclusively in the membrane-associated fraction. This localization could indicate that these proteins might be involved in additional processes such as fatty acid (GlpD) or EPS (GgpP) biosyntheses (Kirsch et al., 2017).

Finally, the salt-induced proteins involved in GG metabolism and transport as well as in G3P synthesis are obviously part of a larger group of salt-induced genes/proteins in *Synechocystis* 6803, which presumably can form a salt-regulon. Using *ggpS* as search string in the CyanoExpress database, which compiles gene expression data sets from *Synechocystis* 6803 (Hernandez-Prieto and Futschik, 2012), revealed more than 30 genes showing similar expression pattern under different environmental stimuli (Suppl. Fig. S6). The occurrence of all genes related to GG biochemistry, which are proven to be involved in salt acclimation, make it very probable that several of the co-regulated genes encode proteins also specific for this stress acclimation. This assumption is supported by our finding that many of these proteins also accumulated to higher levels in salt-acclimated cells. In cases where the protein levels were not significantly elevated, the corresponding gene showed only transient stimulation at the earliest time points after salt addition (Suppl. Fig. S6).

#### Transporters and channels belonging to the salt-specific stimulon

Clustering and functional enrichment analyses clearly indicated that differential regulation of proteins related to membrane transport is generally an important mechanism for salt acclimation (Suppl. Table S9). Among them, the mechanosensitive channel protein of small conductance (MscS) Slr0639 accumulated to the highest level, fold change of 14.3 (RNA approximately 2-fold) (Suppl. Table S6; Fig. 7). Two other MscS proteins, Slr0765 and Sll1040 were identified at elevated protein and RNA levels as well. Msc proteins are important for proper acclimation to hypo-osmotic treatments to facilitate quick release of compatible solutes and to prevent burst of cells (Levina et al., 1999). Hence, similar to elevated GghA, the MscS accumulation likely prepares the cell for upcoming events of lower osmotic pressure as safety valves as has been also shown in many heterotrophic bacteria (Perozo et al., 2001; Stokes et al., 2003). In contrast, the abundance of MscL (Slr0875), which is involved in water movements after sudden osmotic shocks on *Synechocystis* 6803 (Shapiguzew et al., 2005; Azad et al., 2011), did not change in long-term salt-acclimated cells.

**Figure 7.**
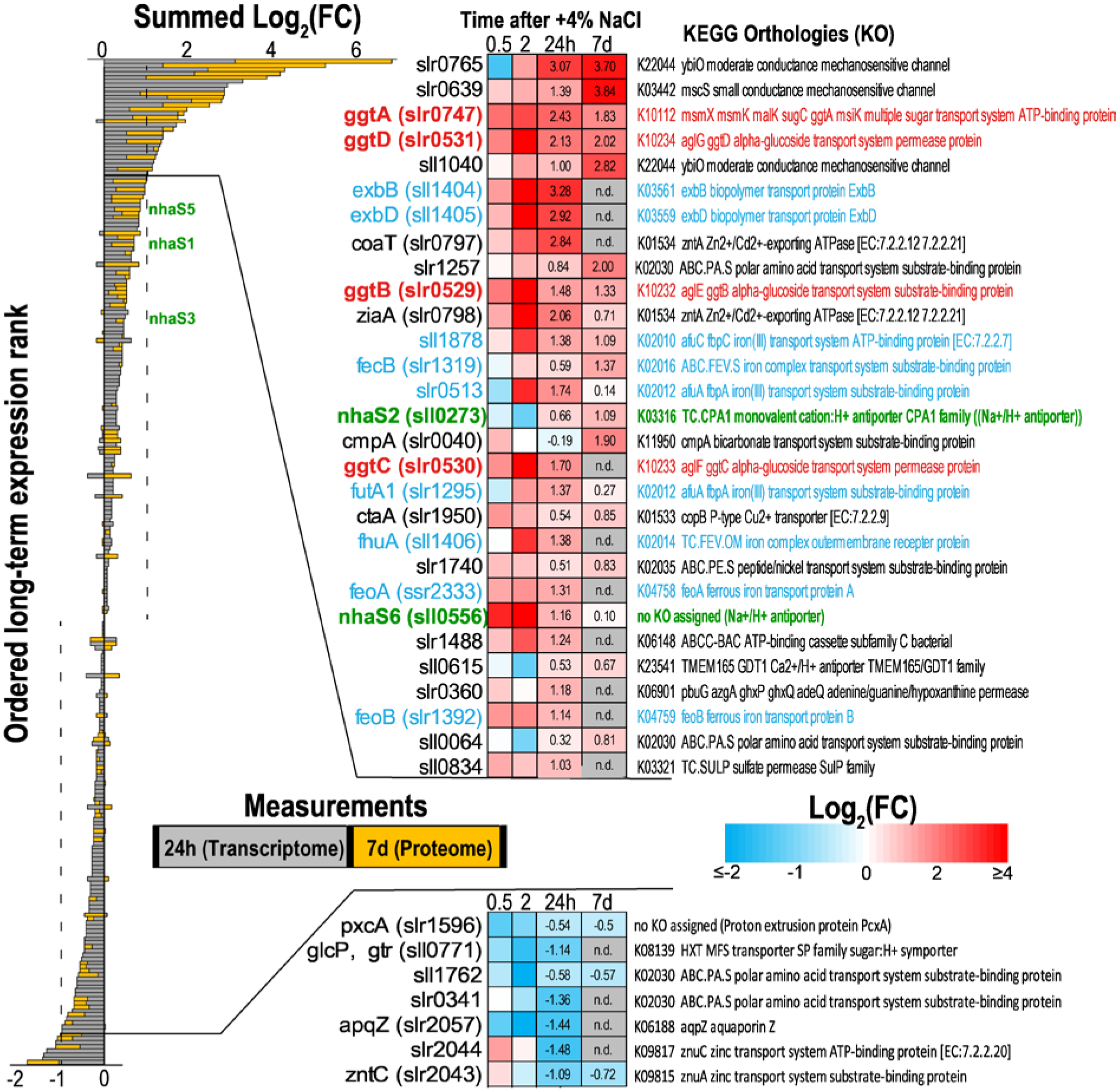
Ordered long-term expression ranks for transport-related genes. Ranks are ordered according to the summed Log_2_(FC) from the 24 h (transcriptome) and 7 d (proteome) measurements of salt-acclimated cells vs. control cells. The heatmaps on the right illustrate the log_2_(FC) at the individual measurements from the microarray experiment (0.5, 2, and 24 h) and proteome measurements (7 d) for the top ranked transport related genes. Highlighted in Red = compatible solute transport (*ggtABCD*); Blue = related to iron transport, Green = *nhaS* genes (Na^+^/H^+^ antiporter). Detailed information is provided in Suppl. Table S9.

Na^+^/H^+^ antiporters are considered as main exporters of excess Na^+^ and, therefore, play a crucial role for salt acclimation (Hagemann, 2011). The *Synechocystis* 6803 genome codes for six Na^+^/H^+^ antiporters, five of them were identified in the proteome. Among them, the protein abundance of NhaS2 and NhaS5 was significantly enhanced in salt-acclimated cells (Fig. 7), whereas the amount of NhaS3, which is essential for cell viability and was discussed to be mainly responsible for Na^+^ export (Wang et al., 2002; Elanskaya et al., 2002), remained unchanged (Suppl. Table S6). However, the fact that the abundance of a protein remains unchanged does not exclude an essential function of this protein for salt acclimation, because it could be regulated on biochemical level according to cellular demands. While crucial roles were assigned to NhaS2 under low Na^+^/K^+^ ratios (Mikkat et al., 2000) or growth at different pH values (Wang et al., 2002), the *nhaS5* mutant did not show any changes compared to wild type (Wang et al., 2002; Elanskaya et al., 2002). Unfortunately, none of the previously discussed candidates for chloride exporters (Sll1864, Slr0753, Sll0855; Hagemann, 2011) could be identified in our proteome data set. Among them, only the gene *sll1864* was transiently stronger expressed on mRNA level in salt-shocked cells (Suppl. Table S1).

In addition, several other proteins potentially involved in ion transport were found in higher abundances in salt-acclimated cells (Fig. 7). For example, two of the 7 annotated cation-transporting ATPases (Wang et al., 2002) were elevated in the proteome (Slr1950 with FC 1.8 and Sll1614 with FC 1.68) and two hours after salt shock in the transcriptome as well. Furthermore, the amount of the protein Slr1257, which comprises a ligand-gated ion channel domain, increased 4-fold, whereas the corresponding mRNA was slightly below the threshold of 2-fold in salt-acclimated cells. Many other transport proteins such as Slr0798 (Zinc-transporting ATPase, FC 1.64), Sll0615 (GDT1-like, possible Ca^2+^/H^+^ antiporter, FC 1.59) or those involved in uptake of nutrients such as nitrate, phosphate, or magnesium were also present in higher amounts in salt-acclimated cells (Suppl. Tables S6, S9). Finally, several transporters involved in iron uptake are transiently up-regulated at mRNA level, which indicates that the transient influx of high NaCl amounts in the cells somehow interferes with the general ion homeostasis including iron availability. However, there expression returned to control mRNA as well as protein levels after long-term salt acclimation (Fig. 7).

#### Many salt-induced proteins are involved in general stress response

Two proteins of unknown function, Sll1862 and sll1863 showed 9.9- and 13.3-fold, respectively, higher abundances in salt-acclimated cells (Table 3). Searches using CyanoExpress (Hernandez-Prieto and Futschik, 2012) revealed that these proteins are induced on mRNA level not only after salt stress but also in response to many other stress treatments, hence, they belong to the group of general stress proteins. The Sll1863 protein was previously identified as the top salt-induced gene/protein (Fulda et al., 2006), while in the present study it shares the top four positions with two MscS proteins, which were not previously quantified. The induction of Sll1862 and Sll1863 also led to high total protein amounts in salt-acclimated cells, both of them belong to the 60 most abundant proteins. However, their inactivation by interposon mutagenesis did not result in a salt-sensitive phenotype (unpublished results of Hagemann group).

Several heat-shock proteins that are involved in protein folding and repair have been previously identified among the salt-induced, general stress proteins (Fulda et al., 2006). In the present study, only the 33 kDa chaperone (Sll1988) was more than 2-fold accumulated while its mRNA showed only a slight increase, whereas the amounts of DnaK, GroEL, or DnaJ proteins were not significantly changed. Moreover, the small, 16.6 kDa heat shock protein (Sll1514) was decreased despite its RNA accumulated after the salt addition (Table 3), whereas it was found before in significant enhanced amounts in salt-acclimated cells (Fulda et al., 2006). These differences most probably result from different cultivation conditions and sampling times in the different studies. Many other salt-stimulated proteins have no functional annotations, however, the proteins Slr0967, Slr2019, Sll0528, and Sll0947 were all also implicated in multiple stress responses of *Synechocystis* 6803 (Uchiyama et al., 2014; Matsuhashi et al., 2015; Lei et al., 2014; Galmozzi et al., 2016).

Moreover, an overlap between salt stress and iron-starvation response has been often observed, because in previous studies many genes/proteins serving as markers for iron-starvation have been observed in elevated amounts in salt-stressed cells as well (Marin et al., 2004; Fulda et al., 2006). In the present study, many proteins related to iron transport are found at higher mRNA levels in the early time points (Fig. 7, Suppl. Table S1). However, in long-term acclimated cells only two iron-regulated proteins are significantly elevated (Table 3). The general stress protein Slr1894, which is annotated as MrgA or Dps like protein, was found in higher abundances in salt-acclimated cells. It has been shown to be involved in oxidative stress response (Li et al., 2004) or in the mobilization of iron-storage after transfer from iron-replete into iron-deplete conditions (Shcolnick et al., 2007). Furthermore, flavodoxin (*isiB*) is clearly accumulated, which plays an important role in the salt-stimulated photosynthetic cyclic electron transport (Hagemann et al., 1999).

#### Salt effects on proteins involved in the basic cell physiology

Previous transcriptome analysis revealed that genes encoding subunits of protein complexes are co-regulated in salt-stressed cells (Marin et al., 2004). The evaluation of the proteome data showed that the abundances of most proteins belonging to the photosystem (PS) I and II, phycobilisomes, ribosomes, or enzymes of the tricarboxylic acid (TCA) cycle were slightly lower after long-term salt acclimation (Fig. 8). Only a few exceptions were found. The Psb28 protein of PSII was elevated (FC 1.8). This subunit is involved in PSII assembly and repair (Nowaczyk et al., 2012), thus its higher amount could indicate that the PSII is less stable in salt-acclimated cells. As another example, the transcript levels of *sll1471* encoding the CpcG2 phycobilisome rod-core linker polypeptide was strongly reduced in the microarray dataset upon short term salt stress and exhibited a lowered protein level (log_2_ FC of −0.42) as well. CpcG2 is the rod core linker of the smaller CpcL-phycobilisome type (Kondo et al., 2005; Liu et al., 2019b), which has been implicated in the formation of a PSI/NDH supercomplex in *Synechocystis* 6803 (Gao et al., 2016). Furthermore, alpha phycocyanobilin lyase (CpcF, *sll1051*), a protein involved in phycobiliprotein assembly into phycobilisomes, increased 1.7-fold. Another phycocyanobilin lyase (CpcE, *slr1878*) was also increased. Finally, many ribosomal proteins showed a tendency to slightly lowered amounts (Fig. 8), which correlates with the slower growth of salt-acclimated cells (Hagemann et al., 1994). The results show that cells acclimated to 684 mM NaCl for 7 days have reached a new steady state, in which many basic physiological processes differ from the control cultures. Regarding basic cellular processes our current transcriptome data of salt-shocked cells after 24 h showed a high agreement with the proteome data.

**Figure 8.**
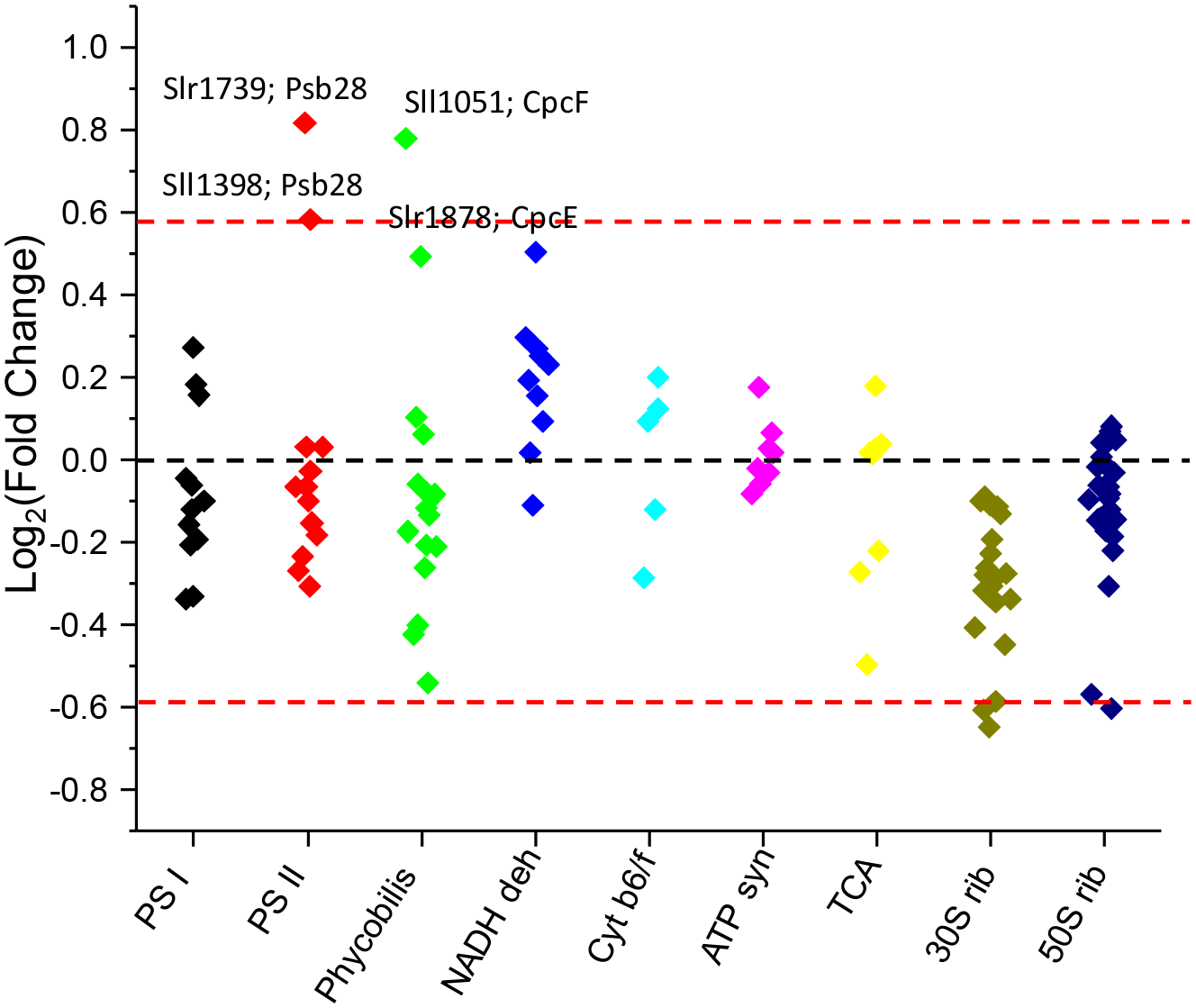
Influence of the salinity on basic cellular processes. Log_2_ fold change valu*e*s (salt-acclimated/control) from the identified protein components of the indicated processes are plotted vertically aligned.

The potential impact of high salinity on the cell surface of *Synechocystis* 6803 cells is indicated by the observation that 8 salt-induced proteins were found among the 19 identified proteins of the functional category „Murein sacculus and peptidoglycan“. These proteins include MurF (Slr1351), DacB (Slr0804), MurA (Slr0017), MurG (slr1656) and MltA (Sll0016), which all are enzymes probably involved in cell wall or cell envelope biogenesis (Suppl. Table S6). Their coordinated up-regulation indicates that salt stress induces a reorganization of cell wall structures, which possibly decrease its permeability for inorganic ions. Changed abundances of several further proteins involved in the reorganization of the cell surface support this assumption. The lowered abundance of Wzc (Sll0923) is consistent with the lowered amount of EPS of salt-acclimated cells (Kirsch et al., 2017). Some of the most giant proteins encoded in the *Synechocystis* 6803 genome that are believed to have functions in the cell envelope, such as the 192 kDa protein Sll0723 (FC 3.9) and the 214 kDa protein Sll1265 (FC 1.6) accumulated in salt-acclimated cells. However, in contrast to the proteins participating in salt-regulated compatible solute synthesis or ion transport, a lower correlation between mRNA and protein abundances was found for the proteins involved cell envelope related processes.

#### Differential expression of regulator proteins

Regulatory proteins are highly important for stress acclimation, but were underrepresented in the proteome data. Only three out of 86 identified regulatory proteins showed significantly elevated protein but not increased mRNA levels in our study. All three represent different members of two-component regulatory systems (Sll1624 – Rre18, Slr1324 – Hik 23, Slr6110 - Rre on plasmid pSYSX). However, none of them is functionally characterized and none of the previously identified salt-stress-associated two-component regulators (Marin et al., 2003; Shoumskaya et al., 2005) showed higher protein amounts here. Further 15 proteins among the 86 identified regulatory proteins were found with lower abundances in salt-acclimated cells, mostly in concordance to their mRNAs.

The list of genes co-regulated with *ggpS* and other genes/proteins for the GG biochemistry included one transcription factor, PrqR (Slr0895; Suppl. Fig. S6). Hence, it is tempting to speculate that PrqR might be involved in the salt-stimulated expression of *ggpS* and other genes in this putative regulon. PrqR was reported as repressor for genes involved in glucose metabolism and oxidative stress acclimation (Khan et al., 2016). The LexA protein, which is not changed on protein level but became significantly decreased on RNA level (Suppl. Fig. S7), has been shown to act as negative regulator of GgpS and GgpP expression (Takashima et al., 2020). However, the fact that the abundance of a protein remains unchanged does not exclude an important function of this protein for salt acclimation, because especially regulatory proteins are often activated/inactivated upon association with other proteins or metabolic signals. Moreover, the expression of genes for other annotated transcription factor changed. For example, the gene *sll0998* encoding the transcription factor RbcR, which regulates the expression of the key CO_2_-fixing enzyme ribulose 1,5 bisphosphate carboxylase/oxygenase (RubisCO), showed lower abundances on protein and RNA levels in salt-acclimated cells. Albeit not passing our significance criteria, both the large and small subunit of RubisCO were reduced by 20-30% that could indeed result in lower CO_2_-fixing activity given the high amount of RubisCO protein per cell.

Finally, sigma factor cascades have been implicated in the stress acclimation of bacteria. Among the down-regulated genes as well as proteins is the anti-sigma F factor antagonist Slr1859 (Suppl. Fig. S7), whose homolog in *Bacillus subtilis* is involved in posttranslational regulation of Sigma factor F (Clarkson et al., 2004). Interestingly, the Sigma F (Slr1564), which itself showed an unchanged protein abundance, is one of the candidate factors involved in mediating the salt-dependent expression of *ggpS* (Marin et al., 2002). Other sigma factors, particularly the genes encoding SigB and SigC were transiently induced after salt addition, while the amount of SigC slightly decreased in the proteome (SigB not identified) (Suppl. Fig. S7). Mutations of these group 2 sigma factors have been shown to reduce the stress tolerance including salt resistance of *Synechocystis* 6803 (Tyystjärvi et al., 2013). Hence, the salt-dependent activity of different sigma factors might significantly contribute to the observed salt-induced expression changes on transcript and protein levels.

#### Salt stress differentially affects chromosome regions

Most salt-regulated proteins are encoded by single genes or in small operons that are spread on the *Synechocystis* 6803 chromosome. However, some salt-regulated genes are clustered on specific chromosomal regions. The most remarkable examples are clusters including the genes *ggpS* and *ggpP*, which were found in regions on the chromosome comprising several genes/proteins with salt-stimulated expression. The *ggpS* cluster comprises 9 salt-regulated genes, which are at least transiently stronger expressed in salt-grown cells than in control cells (Suppl. Fig. S8). Upstream of the *ggpS/glpD* operon, on the opposite strand, a salt-regulated gene cluster of at least 7 genes (*slr1670-1677*) is found. This gene organization is widely conserved in the genomes of many GG-accumulating cyanobacteria (Kirsch et al., 2017). A second salt-stimulated gene cluster can be found downstream of *ggpP* and *ggtA*, which comprises 5 different genes (Suppl. Fig. S9). In addition to *glpK* four other genes (three encode subunits of protochlorophyllide synthase, e.g. ChlN, *slr0750*; see Table 3) are at least transiently salt-regulated but have not yet shown to code for proteins directly involved in salt acclimation.

Another salt-stimulated gene cluster was found on the plasmid pSYSA. The genes *sll7063-sll7067* (for one example see Table 3) are *cas* genes that encode structural proteins of one of the three CRISPR-Cas systems in *Synechocystis* 6803, called CRISPR2 (Scholz et al., 2013). The five proteins were significantly up-regulated whereas the transcripts were transiently down-regulated and then slightly up-regulated after 24 h of salt acclimation. In contrast, the CRISPR3 system (*sll7085-sll7090*) on the same plasmid was consistently down-regulated at RNA and protein level. Recently, it has been shown that these proteins form a stable protein complex together with their cognate crRNAs (Riediger et al., 2021).

There are also some examples of coordinately down-regulated genes/proteins in salt-acclimated cells. One example is the large region (chromosome positions 1181250 – 1200000) comprising 22 genes, which are forming three operons (Suppl. Fig. S10). The first operon *slr1406-1410* encodes proteins of unknown function. The second salt-repressed operon is situated on the opposite strand (*sll1307-1304* and *sll1784-1785*) and also mostly encodes not functionally annotated proteins; however, the protein Sll1305 resembles ketose 3-epimerases while Sll1307 and Sll1784 are predicted outer membrane-bound periplasmic proteins. Hence it can be speculated that these proteins are somehow involved in cell wall synthesis/reorganization. The third salt-repressed operon in this region comprises the genes *slr1852-1862*. This cluster contains several annotated genes (Slr1855 N-acetylglucosamine 2-epimerase, Slr1856 – anti-sigma factor antagonist, GlgX1 – glycogen branching enzyme 1, IcfG – carbon metabolism regulator), which play important roles in the primary carbon metabolism and, particularly, its regulation (Beuf et al., 1994; Shi et al., 1999). Another example of coordinated down-regulated genes/proteins represents the large operon *slr0144-0152*, which has been noted before as one of the highly coordinated expressed regions on the chromosome of *Synechocystis* 6803 (e.g., Summerfield and Sherman, 2008) that is controlled by the redox-responsive transcription factor RpaB (Riediger et al., 2019).

#### Metabolome analysis of salt-acclimated cells

The presence of high NaCl amounts in the medium induces a massive GG accumulation (Fig. 6C), which likely triggers a strong redistribution of organic carbon in *Synechocystis* 6803. The large impact of GG synthesis on overall carbon metabolism is also consistent with the observation that many proteins (and their genes) involved in glycogen metabolism as well as glycolysis showed significant changes in their abundances (Table 4). For example, the neopullulanase, glycogen phosphorylase GlgP2 (Slr1367) and one debranching enzyme GlgX1 (Slr0237, the other one Sll1857 is decreased) showed higher abundances on protein as well as RNA levels in salt-acclimated cells. The different response of the two GlgX proteins towards salt stress has been previously shown with Western-blotting (Iijima et al., 2015). Hence, the demand of organic carbon for the synthesis of GG precursors is at least partly supported by an enhanced glycogen breakdown and reduced glycogen build up, because glycogen and GG synthesis are competing for the same precursor, ADP glucose. The relatively low carbon/nitrogen state in salt-acclimated cells is also reflected by the lowered amount of 2-oxoglutarate (2OG, Fig. 9), which is the key metabolic signal reporting changes of the cellular carbon/nitrogen ratio in cyanobacteria (Hagemann et al., 2021).

**Figure 9.**
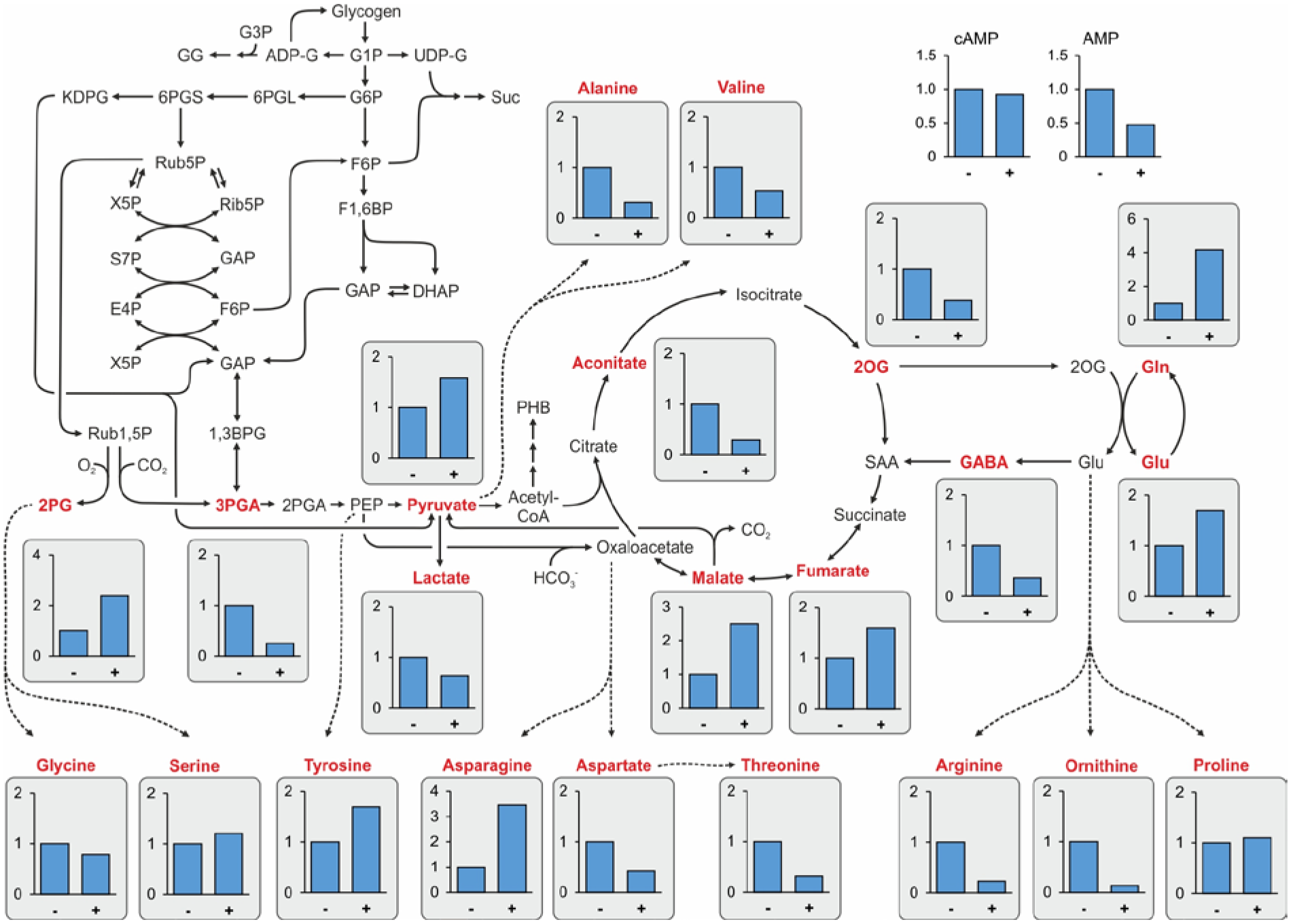
Alterations in the central carbon and nitrogen metabolism. Low molecular mass compounds were isolated from cells of *Synechocystis* 6803 grown in NaCl-free BG11 medium or medium supplemented with 4% NaCl for 7 days. LC-MS/MS was used to estimate the relative levels (Y axis show fold changes, amount in cells from 0% NaCl cultivation (-) set to 1 and relative level at 4% NaCl (+) is shown) of central metabolites as part of primary carbon and nitrogen metabolism. Shown are mean values from three biological replicates (details in Suppl. Table S10). Dotted arrows indicate that several enzymes are necessary to convert one metabolite into the other. The alteration in the compatible solute GG is shown in Fig. 6.

To obtain a snapshot on metabolites of the central carbon and nitrogen metabolism, LC-MS/MS was used (Suppl. Table S10). The relative levels of the RubisCO carboxylation and oxygenation products 3PGA and 2PG, respectively, showed opposite behavior (Fig. 9). 3PGA accumulated approximately 3-fold less in salt-acclimated cells, while 2PG was clearly enhanced. This could indicate a decreased CO_2_/O_2_ ratio at the active site of Rubisco, since the amounts of these gasses mainly regulate its relative carboxylation/oxygenation activity. For example, it might be possible that due to the higher content of inorganic ions inside the salt-exposed cells carboxysomes are less gas tight in high salt-grown cells, thereby promoting a better diffusion of oxygen into carboxysomes reducing the CO_2_/O_2_ ratio. The observed changes in the 3PGA and 2PG levels are consistent with the reported lower photosynthetic activity and growth rate in salt-acclimated cells of *Synechocystis* 6803 (e.g., Hagemann et al., 1994), which certainly also reduce Calvin-Benson-cycle activity. Consistently, the protein abundances of photosynthetic complexes, Calvin-Benson-cycle enzymes including RubisCO and components of the cyanobacterial inorganic carbon-concentrating mechanism were found at 10-40% lower levels, which is below our significance threshold but might contribute to lower photosynthetic activity in salt-acclimated cells. Furthermore, more organic carbon could be taken out from the Calvin-Benson-cycle, which is for example seen in the increased amount of pyruvate and organic acids in the reductive branch of the TCA cycle, such as malate and fumarate. This interpretation is also supported by the finding that Gap1, the glyceraldehyde dehydrogenase 1 (Slr0884) involved in sugar catabolism (Koksharova et al., 1998) is also up-regulated (Table 4), while Gap2 involved in photosynthetic carbon assimilation did not change (Suppl. Table S1).

In addition to the carbon fixation and allocation, nitrogen assimilation is altered in salt-acclimated cells, which is reflected by enhanced glutamine and glutamate levels while 2OG, the carbon skeleton used for ammonia assimilation decreased (Fig. 9). Increased glutamate levels have been often reported in salt-exposed bacteria (Hagemann, 2011), because this negatively charged amino acid is compensating the positive charge of cations, especially K^+^. The enzymes involved in the GS/GOGAT cycle for assimilation of NH_4_^+^ into 2OG did not significantly change their expression in long-term salt acclimated cells, but were significantly lowered immediately after salt addition. However, the glutamate decarboxylase Gad (Sll1641) showed increased expression in salt-grown cells. Proline, which is often used in heterotrophic bacteria as compatible solute and can be synthesized from glutamate, is not changed during salt acclimation of *Synechocystis* 6803 (Fig. 9). Moreover, aspartate and arginine levels decreased. These amino acids serve as precursors for cyanophycin, the nitrogen storage compound of *Synechocystis* 6803. In this regard it is interesting to note that cyanophycin synthetase (Slr2002, FC 0.64) was decreased on protein level while the mRNA did not change. Furthermore, the mutation of a gene presumably involved in cyanophycin turnover resulted in a salt-sensitive phenotype of *Synechocystis* 6803 (Zuther et al., 1998). These observations also add to the assumption of altered nitrogen assimilation in salt-loaded cells. Marked changes in the amino acid composition had been also reported for *Synechocystis* 6803 cells when grown in artificial seawater medium compared to BG11 (Iijima et al., 2015). Similarly, global changes in the carbon- and nitrogen metabolism have been noticed in *Synechococcus* sp. PCC 7002 (Aikawa et al., 2019).

## Discussion

### Overall correlation of omics data

Our proteomic and transcriptomic data showed a good relationship (Pearson correlation coefficient, 0.58) especially taking into consideration that sampling was done in different laboratories and at different time periods after salt treatment. This finding indicates that in most cases transcriptional activation/repression leads to enhanced/diminished protein amounts, whereas the differential regulated ncRNAs rather regulate single genes/proteins. Generally, relationships between the transcript and corresponding protein amounts are influenced by several processes, such as (1) impact of mRNA processing or ncRNAs on translation rates, (2) proteińs half-life, and (3) protein synthesis delay (Liu et al., 2016). Other studies found widely differing correlation coefficients between cyanobacterial transcriptomes and proteomes depending on the conditions examined. For example, experiments with *Synechocystis* 6803 cells shifted from high to low CO_2_ conditions showed also a good correspondence between transcriptomics and proteomics (Spät et al., 2021), whereas Toyoshima et al. (2020) reported a rather low correlation between transcriptomics and proteomics in *Synechocystis* 6803 cells grown under phototrophic, mixotrophic or heterotrophic conditions. Similarly, a very low correlation was reported from an integrated proteomic and transcriptomic analysis of salt stress responses in *Synechocystis* 6803 (Qiao et al., 2013).

Moreover, the comparison of proteome and transcriptome data from different studies, despite the varying degree of correspondence, offers the possibility to filter out regularly, truly salt-regulated proteins. To this end, we compared three transcriptomic with our proteomic data sets to identify further proteins potentially involved with unknown function in salt acclimation. Filtering the four datasets for features that were at least 1.5-fold increased (log_2_ > 0.58) in the three transcriptome data sets and at least 1.3-fold increased (log_2_ > 0.38) in the proteome data set produced a list of 44 genes, for which in 25 cases the corresponding proteins were identified (Suppl. Table S11). This group of genes/proteins includes 11 that are involved in GG metabolism and ion transport. Among the rather general stress proteins with unknown function are the Sll1862 and Sll1863 proteins as well as the putative zinc metalloprotease Sll0528 and the Slr1894 protein, which all have been discussed before. Other proteins with unknown function in salt acclimation such as the methionine aminopeptidase B (MAP B, Slr0786; co-expressed with *ggpS*, Suppl. Fig. S6), Sll0723 and Slr0001 are candidate proteins for further studies. Among the 19 salt-induced genes, whose proteins were not identified in our study, only the *slr0530* gene product as component of the GG transporter is functionally related to known processes in salt acclimation.

The good correlation of transcriptomic and proteomic changes extends to the alterations on the metabolome level. Basic alterations in the central carbon and nitrogen metabolism are also supported by expression changes. However, biochemical alteration due to changed metabolite and ion levels on key enzyme activities certainly contribute to the novel metabolic homeostasis in salt-acclimated cells as has been recently discussed for the metabolic acclimation of *Synechocystis* 6803 towards different CO_2_ availability (Jablonsky et al., 2016).

### Salt shock leads to a temporally staggered reshaping of the transcriptome composition

The salt acclimation response goes way beyond the induction of gene expression required for the compatible solute machinery, it has clearly a great impact on general metabolism. Along the temporal axis, the reprogramming of gene expression can be differentiated between an early response with the respective minima and maxima leading to the four different clusters (Fig. 2). The metabolic response included the rapid repression of the ammonia assimilation system, detected by decreased transcript abundances for *amt1* and *glnA* encoding the ammonium transporter and the primary enzyme for ammonium incorporation, glutamine synthetase (GS). Consistently, increased transcript levels were found for genes *gifA* and *gifB*, which encode inhibitory proteins for GS, thereby blocking ammonium assimilation (for an overview see Bolay et al., 2018). After 24 h the transcript levels of these genes were at the initial levels.

The question arises, which processes are responsible for the staggered reshaping of the transcriptome. It has been shown that shortly after salt addition the cytoplasmic composition underwent rapid changes due to ion and water movements, whereby the early high internal ion contents, especially of Na^+^ are discussed to inhibit metabolic activities but also to trigger acclimation responses such as GG synthesis activation (reviewed in Hagemann, 2011). The transporters responsible for the rapid ion movements are largely unknown, especially verified candidates for Cl^-^ export are still missing. In the present study we did not find marked expression changes for genes encoding potential anion exporters. It can be assumed that these transporters are mainly regulated on their activity levels to manage the ion regulation within the first minutes to hours after salt shock, because *de novo* protein synthesis is one process clearly down-regulated after salt shock (Hagemann et al., 1994). However, we found that especially in the early time points after salt shock multiple ncRNAs become up- or down-regulated, which were not covered in the previous transcriptomic datasets. These small RNAs likely reshape mainly the translational efficiency of specific target mRNAs. Hence, it is well possible that many of the identified asRNAs and sRNAs are important to fine tune the translational response in the acute stress situation. Further work is necessary to identify the specific targets of the sRNAs and to verify the action of the ncRNAs during salt acclimation in cyanobacteria.

### Impact of salt on DNA structure

The identification of chromosomal regions in the *Synechocystis* 6803 genome with coordinately up- or down-regulated genes under high salt conditions indicates that the DNA structure likely differs between distinct chromosomal regions (Suppl. Figs. S8-10). Similar observations were made when the impact of antibiotics that affect DNA supercoiling was analyzed on global gene expression patterns in *E. coli* and *Synechocystis* 6803. Interestingly, a large overlap has been observed between the antibiotic-induced gene expression changes and the salt and osmostress responses (Cheung et al., 2003; Prakash et al., 2009). Hence, the salt-stimulated expression of genes in some chromosomal regions could be related to a relaxed DNA topology permitting easier access of the transcription machinery and *vice versa*. In this regard it is interesting to note that the gene *slr2058* encoding the DNA topisomerase I is higher transcribed in the first hours after salt shock (Suppl. Tables S1 and S2). The ATP-dependent DNA topoisomerases relax negative supercoils and are specifically involved in chromosome partitioning, which has been shown to be of fundamental importance for bacterial gene expression (Dorman and Dorman, 2016). Recently, another subunit of DNA toposiomerase has been suggested to be involved in the regulation of DNA replication in a mutant defective in one of the dominating DNA methylation activities in *Synechocystis* 6803 (Gärtner et al., 2019). It might be possible that differences in DNA methylation and thereby induced changes in the DNA/protein association are at least partly responsible for the different accessibility of specific chromosomal regions under different salt conditions. Finally, the interaction of chromosomal DNA with the GG synthesizing enzyme GgpS has been shown to be central for the ion-mediated activation/inactivation of its biochemical activity (Novak et al., 2011). Collectively, our data support the notion that changes in DNA structure and DNA/protein interactions due to altered ionic and electrostatic relations play an important role in microbial salt acclimation.

### Future developments – regulation and application

One still open question is how salt stress is sensed and transduced to the cellular gene expression machinery. Despite several efforts, specific salt-sensing proteins have not been identified in cyanobacteria. The screening of mutant collections defective in histidine kinases and cooperating response regulators identified some two-component systems that are involved in the salt-induced regulation of different general stress proteins, however, the induction of salt-specific proteins including *ggpS* remained unchanged (Marin et al., 2003; Shoumskaya et al., 2005). Two proteins were identified as repressors for *ggpS*, the small GgpR protein (Klähn et al., 2010b) and the transcriptional factor LexA have been shown to bind specifically the *ggpS* promoter (Takashima et al., 2020). However, the role of LexA as specific salt-sensing transcription factor is unlikely, because it is also involved in the regulation of many other processes in *Synechocystis* 6803, for example a verified role in fatty acid accumulation (Kizawa et al., 2017), and the *lexA* mutant has no reported salt-sensitive phenotype or changed GG accumulation. The present study identified two other likely candidates for a salt-stress specific gene regulation. First, the transcription factor PrqR (Slr0895) represents an interesting candidate, because the gene *sl0895* is clearly co-regulated with many genes coding proteins involved on GG metabolism (Suppl. Fig. S6). It has been recently shown that PrqR is involved in the acclimation to oxidative stress in *Synechocystis* 6803 (Khan et al., 2016). Salt stress is also inducing oxidative stress in cyanobacterial cells, hence, the finding of the role of PrqR in this stress acclimation process might of secondary importance. Second, the gene *ssl1326* that possibly encodes a CopG family transcription factor has been found strongly induced after salt shock in the DNA microarray data set (Suppl. Table S1). Further work is necessary to validate whether PrqR or CopG are somehow acting as salt-specific gene expression regulators. In the moment, it might well be possible that ion-mediated changes in the DNA structure, RNA-polymerase affinity, and enzyme activities might be the main and sufficient mechanism to acclimate towards different salt conditions in euryhaline bacteria such as *Synechocystis* 6803.

Furthermore, salt acclimation is becoming more important for applied research with cyanobacteria regarding the direct use of compatible solutes as well as mass cultivation in sea water to make the process more sustainable (Pade and Hagemann, 2014; Cui et al., 2020). For example, the cyanobacterial production of mannitol (Wu et al., 2020) and trehalose (Qiao et al., 2020) has been promoted by cultivation at enhanced salinities. Moreover, a more salt-tolerant version of the fast-growing *Synechococcus elongatus* strain UTEX2973 has been engineered by the expression of GG synthesis genes, which can be used for biotechnological purposes in full marine waters (Cui et al., 2021) However, saline conditions might also negatively affect the production titer. The large impact of GG accumulation on the overall carbon metabolism also negatively influenced the ethanol production with *Synechocystis* 6803 at 4% NaCl while 2% NaCl were slightly stimulatory (Pade et al., 2017). Salt-regulated, strong promoters might be an option to improve transgene expression in salt-grown cyanobacteria. Hence, a deeper knowledge on salt acclimation will promote future biotechnological applications with cyanobacteria.

## Material and Methods

### Cultivation and sampling

*Synechocystis* sp. 6803 substrain PCC-M was used in all experiments. Axenic cells were maintained on agar plates with BG11 mineral medium at 30°C under constant illumination. For salt stress experiments, axenic cells were grown photoautotrophically in glass tubes containing liquid BG11 medium (TES-buffered at pH 8) at 29°C in the cooperating laboratories. If not stated differently, cultures were aerated with CO_2_-enriched air (5% CO_2_ [v/v]) and kept under continuous illumination of 150 µmol photons m^-2^ s^-1^ (warm light, Osram L58 W32/3). Control cells were cultured in NaCl-free BG11 medium, whereas salt-acclimated cultures were obtained after long-term growth (up to one week) in BG11 medium supplemented with 684 mM NaCl (4% NaCl [w/v]). For this, cells were transferred daily to fresh media with 4% NaCl. Short-term salt shock experiments were performed by adding crystalline NaCl to a final amount of 4% into control cultures at time point zero. Subsequently, cells were harvested at defined time points.

For proteomics, cells were harvested by centrifugation at 14,000 g and 4°C for 5 min. The cell pellets were frozen in liquid nitrogen and stored at −80°C. For metabolomics, cells were harvested by quick filtration on nitrocellulose filters (0,45 µm pore size) in growth light within 30 s. Cells on filters were then frozen in liquid nitrogen and stored at −80°C. For transcriptomics, cells from 40 ml culture were harvested by quick filtration on hydrophilic polyethersulfone filters (Pall Supor-800, 0.8 µm), immediately immersed in 1 ml of cold PGTX solution (Pinto et al., 2009) and frozen in liquid nitrogen.

### Transcriptomic methods

#### RNA extraction

Total RNA was extracted as described previously (Hein et al., 2013). Prior to the microarray analysis, 10 µg of total RNA were treated with Turbo DNase (Invitrogen) according to the manufacturer’s protocol and precipitated with ethanol/sodium acetate. RNA quality was verified by electrophoresis on MEN-buffered 1.5% agarose gels supplemented with 6% formaldehyde.

#### Microarray analysis

To enable the comparison with a previous microarray analysis (Marin et al., 2004), we analyzed similar time points after salt shock: 0, 0.5, 2 and 24 h. Labeling and hybridization were performed as described (Klähn et al., 2015). Three µg of RNA were used for the labeling reaction and 1.65 µg of labeled RNA for the hybridization. The microarray hybridization was performed in duplicates for each sampling point. Almost all features on the microarray chip were covered by several independent probes. In addition, it contained technical replicates for each single probe. Hence, mean values for all probes of a given feature were used for the final calculation of fold changes. Data processing and statistical evaluation was performed using the R software as described (Klähn et al., 2015). The full dataset is accessible from the GEO database with the accession number GSE174316 (accession for Reviewers via: izupwswenzelfkv).

In Suppl. Table S1 transcripts are separated into mRNAs, asRNAs, other ncRNAs, 5’UTRs and transcripts derived from internal TSSs, i.e. within CDS (int). However, it should be noted that for every category overlaps are possible, i.e. the annotation as well as microarray detection of ncRNAs is often ambiguous since they can overlap with UTRs. Thus, all features labeled with the systematic term “NC-#” are referred as “potentially” trans-acting ncRNAs. For the evaluation presented in the text only ncRNAs with an annotation based on Mitschke et al. (2011) were considered.

#### Cluster analysis

Clustering analysis for the microarray time series was performed using the *mfuzz* R package (Kumar and Futschik, 2007). To reduce noise and false positives, only 3831 transcripts entered the clustering analysis that had a difference in their absolute expression values higher than log_2_ ≥│1│in at least one of these comparisons. The optimal cluster number was determined with the Elbow method using the minimum centroid distance and an estimated “*fuzzifier”* parameter of m = 2.53, yielding four clusters. Transcripts with low membership values < 0.5 were removed. However, those transcripts were still included if the combined membership values for two similar clusters (e.g. cluster 1 + 2 and 3 + 4) exceeded > 0.6.

### Proteomic methods

Cells from four biological replicates of salt-acclimated and control cultures, respectively, were broken with glass beads using Precellys 24 homogenizer (peqLab Biotechnologie GmbH, Erlangen, Germany) in non-denaturing buffer containing 10 mM Tris/HCl, pH 7.4, 138 mM NaCl, 2.7 mM KCl, 1 mM MgCl_2_. After withdrawing an aliquot of each cell extract as total protein the remaining cell extracts were fractionated into debris, soluble, and membrane-enriched fractions. Protein samples were reduced with dithiothreitol, alkylated with iodoacetamide and digested with trypsin in sodium deoxycholate-containing buffer solution (Pappesch et al. 2017).

LC-HDMS^E^ analyses of desalted peptide samples supplemented with 40 fmol of Hi3 Phos B standard for protein absolute quantification (Waters) were carried out using a nanoAcquity UPLC system (Waters) coupled to a Waters Synapt G2-S mass spectrometer (Pade et al., 2017). The Synapt G2-S instrument was operated in data-independent mode with ion-mobility separation as an additional dimension of separation (referred to as HDMS^E^). Single measurements of the four biological replicates of the total extract, the soluble fraction and the membrane-enriched fraction were carried out, while pooled samples of the debris fraction were measured in triplicate.

Progenesis QI for Proteomics version 4.1 (Nonlinear Dynamics, Newcastle upon Tyne, UK) was used for raw data processing, protein identification and label free quantification. Proteins were quantified by the absolute quantification Hi3 method using Hi3 Phos B Standard (Waters) as reference (Silva et al. 2006). Results were given as fmol on column. Proteins identified in one fraction by at least two unique peptides were included in the quantitative analysis. Protein abundance changes between salt-acclimated and control cells by a factor of at least 1.5, accompanied by ANOVA *p*-values < 0.05 were regarded as significant. To determine a combined fold change value for the total extract and the three subcellular fractions, first a weighted fold change value for the three subcellular fractions was calculated by summing up the corresponding protein amounts (measured as fmol on column) and dividing the value from salt-acclimated cells by the value from controls. Since the debris fraction contained only a minor part of about 10% of the total protein, the protein amounts of the debris fraction were divided by ten. In a second step, the final combined fold change value was calculated as the average of the weighted fold change from the subcellular fractions and the fold change of the total extract.

A detailed description of the proteome method can be found in the Supplementary Material. The mass spectrometry proteomics data have been deposited to the ProteomeXchange Consortium via the PRIDE (Vizcaíno et al., 2013) partner repository with the dataset identifier PXD026118 and 10.6019/PXD026118. Data are available via ProteomeXchange with identifier PXD026118 (Reviewer account details: Username; Password: NjhSNM7Z).

### Metabolomic methods

Low molecular mass compounds were extracted from cells with ethanol (80%, HPLC grade, Roth, Germany) at 65°C for 2 h. The soluble sugars were analyzed by gas-liquid chromatography using a defined amount of sorbitol as internal standard as described by Hagemann et al. (2008). Other metabolites were analyzed by LC-MS/MS using the high-performance liquid chromatograph mass spectrometer system LCMS-8050 (Shimadzu, Japan). One microgram of carnitine was added per sample as internal standard for LC-MS/MS analyses. The dry extracts were dissolved in 200 µl MS-grade water and filtered through 0.2 µm filters (Omnifix®-F, Braun, Germany). The cleared supernatants were separated on a pentafluorophenylpropyl (PFPP) column (Supelco Discovery HS FS, 3 µm, 150 x 2.1 mm) with a mobile phase containing 0.1% formic acid. The compounds were eluted at a rate of 0.25 ml min^-1^ using the following gradient: 1 min 0.1% formic acid, 95% water, 5% acetonitrile, within 15 min linear gradient to 0.1% formic acid, 5% distilled water, 95% acetonitrile, 10 min 0.1% formic acid, 5% distilled water, 95% acetonitrile. Aliquots were continuously injected in the MS/MS part and ionized via electrospray ionization (ESI). The compounds were identified and quantified using the multiple reaction monitoring (MRM) values given in the LC-MS/MS method package and the LabSolutions software package (Shimadzu, Japan). The metabolites were determined as relative metabolite abundances (fold changes), which were calculated after normalization of signal intensity to that of the internal standard carnitine.

## Supporting information

Suppl. Genome plot

Suppl. Data sets

## ACKNOWLEDGEMENTS

The technical assistance of Klaudia Michl (University of Rostock) is acknowledged.

## AUTHOR CONTRIBUTION

MH designed the study. SK, MR, JG & WRH performed and evaluated the transcriptomic analyses. SM performed and evaluated the proteome experiments. MH performed and evaluated the metabolome analyses. MH, WRH, SK & SM wrote the manuscript.

## References

Aikawa S, Nishida A, Hasunuma T, Chang JS, Kondo A (2019) Short-term temporal metabolic behavior in halophilic cyanobacterium *Synechococcus* sp. strain PCC 7002 after salt shock. Metabolites 9: 297

Azad AK, Sato R, Ohtani K, Sawa Y, Ishikawa T, Shibata H (2011) Functional characterization and hyperosmotic regulation of aquaporin in *Synechocystis* sp. PCC 6803. Plant Sci 180: 375–382

Beuf L, Bédu S, Durand MC, Joset F (1994) A protein involved in co-ordinated regulation of inorganic carbon and glucose metabolism in the facultative photoautotrophic cyanobacterium *Synechocystis* PCC6803. Plant Mol Biol 25: 855–864

Billis K, Billini M, Tripp HJ, Kyrpides NC, Mavromatis K. 2014. Comparative transcriptomics between *Synechococcus* PCC 7942 and *Synechocystis* PCC 6803 provide insights into mechanisms of stress acclimation. PLoS One 9: e109738.

Bolay P, Muro-Pastor MI, Francisco J Florencio FJ, Klähn S (2018) The distinctive regulation of cyanobacterial glutamine synthetase. Life 8: 52

Cheung KJ, Badarinarayana V, Selinger DW, Janse D, Church GM (2003) A microarray-based antibiotic screen identifies a regulatory role for supercoiling in the osmotic stress response of *Escherichia coli*. Genome Res 13: 206–215

Chisti Y (2013) Constraints to commercialization of algal fuels. J Biotech 167: 201–214

Clarkson J, Campbell ID, Yudkin MD (2004) Efficient regulation of sigmaF, the first sporulation-specific sigma factor in *B. subtilis*. J Mol Biol 342: 1187–1195

Cui J, Sun T, Chen L, Zhang W (2020) Engineering salt tolerance of photosynthetic cyanobacteria for seawater utilization. Biotechnol Adv 43: 107578

Cui J, Sun T, Chen L, Zhang W (2021) Salt-tolerant *Synechococcus elongatus* UTEX 2973 obtained via engineering of heterologous synthesis of compatible solute glucosylglycerol. Front Microbiol 12: 650217

Dorman CJ, Dorman MJ (2016) DNA supercoiling is a fundamental regulatory principle in the control of bacterial gene expression. Biophys Rev 8: 209–220

Dühring U, Axmann IM, Hess WR, Wilde A (2006) An internal antisense RNA regulates expression of the photosynthesis gene isiA. Proc Natl Acad Sci USA 103: 7054–7058

Eisenhut M, Georg J, Klähn S, Sakurai I, Mustila H, Zhang P, Hess WR, Aro EM (2012) The antisense RNA As1_flv4 in the cyanobacterium *Synechocystis* sp. PCC 6803 prevents premature expression of the flv4-2 operon upon shift in inorganic carbon supply. J Biol Chem 287: 33153–33162

Elanskaya IV, Karandashova IV, Bogachev AV, Hagemann M (2002) Functional analysis of the Na^+^/H^+^ antiporter encoding genes of the cyanobacterium Synechocystis PCC 6803. Biochemistry (Moscow) 67: 432–440

Fulda S, Mikkat S, Huang F, Huckauf J, Marin K, Norling B, Hagemann M (2006) Proteome analysis of salt stress response in the cyanobacterium Synechocystis sp. strain PCC 6803. Proteomics 6: 2733–2745

Galmozzi CV, Florencio FJ, Muro-Pastor MI (2016) The cyanobacterial ribosomal-associated protein LrtA is involved in post-stress survival in *Synechocystis* sp PCC 6803. PLoS One 11: e0159346

Gao L, Wang J, Ge H, Fang L, Zhang Y, Huang X, Wang Y (2015a)Toward the complete proteome of *Synechocystis* sp. PCC 6803. Photosynth Res 126: 203–219

Gao L, Ge H, Huang X, Liu K, Zhang Y, Xu W, Wang Y (2015b) Systematically ranking the tightness of membrane association for peripheral membrane proteins (PMPs). Mol Cell Proteomics 14: 340–353

Gao F, Zhao J, Chen L, Battchikova N, Ran Z, Aro EM, Ogawa T, Ma W (2016) The NDH-1L-PSI supercomplex Is important for efficient cyclic electron transport in cyanobacteria. Plant Phys 172: 1451–1464

Georg J, Dienst D, Schürgers N, Wallner T, Kopp D, Stazic D, et al. (2014) The small regulatory RNA SyR1/PsrR1 controls photosynthetic functions in cyanobacteria. Plant Cell 26: 3661–3679

Georg J, Kostova G, Vurijoki L, Schön V, Kadowaki T, Huokko T, et al. 2017. Acclimation of oxygenic photosynthesis to iron starvation is controlled by the sRNA IsaR1. Curr Biol 27: 1425–1436

Georg J, Hess WR (2018) Wide-spread antisense transcription in prokaryotes. Microbiol Spectrum 6: RWR–0029-2018

Grigorieva G, Shestakov S. 1982. Transformation in the cyanobacterium *Synechocystis* sp. 6803. FEMS Microbiol Lett 13: 367–370

Gärtner K, Klähn S, Watanabe S, Mikkat S, Scholz I, Hess WR, Hagemann M (2019) Cytosine N4-methylation via M.Ssp6803II is involved in the regulation of transcription, fine-tuning of DNA replication and DNA repair in the cyanobacterium *Synechocystis* sp. PCC 6803. Front Microbiol 10: 1233

Hagemann M, Fulda S, Schubert H (1994) DNA, RNA and protein synthesis in the cyanobacterium *Synechocystis* sp. PCC 6803 adapted to different salt concentrations. Curr Microbiol 28: 201–207

Hagemann M, Schoor A, Jeanjean R, Zuther E, Joset F (1997) The *stpA* gene form *Synechocystis* sp. strain PCC 6803 encodes the glucosylglycerol-phosphate phosphatase involved in cyanobacterial osmotic response to salt shock. J Bacteriol 179: 1727–1733

Hagemann M, Jeanjean R, Fulda S, Havaux M, Erdmann N (1999) Flavodoxin accumulation contributes to enhanced cyclic electron flow around photosystem I in salt-stressed cells of *Synechocystis* sp. PCC 6803. Physiol Plant 105: 670–678

Hagemann M, Ribbeck-Busch K, Klähn S, Hasse D, Steinbruch R, Berg G (2008) The plant-associated bacterium *Stenotrophomonas rhizophila* expresses a new enzyme for the synthesis of the compatible solute glucosylglycerol. J Bacteriol 190: 5898–5906

Hagemann M (2011) Molecular biology of cyanobacterial salt acclimation. FEMS Microbiol Rev 35: 87–123

Hagemann M, Hess WR (2018) Systems and synthetic biology for the biotechnological application of cyanobacteria. Curr Opin Biotech 49: 94–99

Hagemann M, Song S, Brouwer EM (2021) Inorganic carbon assimilation in cyanobacteria: Mechanisms, regulation, and engineering. In Hudson P, Lee SY, Nielsen J (eds.) Cyanobacteria Biotechnology, Wiley-Blackwell Biotechnology Series, Chapter 1, 1–31.

Hernandez-Prieto MA, Futschik ME (2012) CyanoEXpress: A web database for exploration and visualisation of the integrated transcriptome of cyanobacterium *Synechocystis* sp. PCC6803. Bioinformation 8: 634–638

Hein, S., Scholz, I., Voß, B., and Hess, W. R. (2013). Adaptation and modification of three CRISPR loci in two closely related cyanobacteria. RNA Biol 10: 852–864

Huang F, Fulda S, Hagemann M, Norling B (2006) Proteomic screening of salt-stress-induced changes in plasma membranes of *Synechocystis* sp. strain PCC 6803. Proteomics 6: 910–920

Iijima H, Nakaya Y, Kuwahara A, Hirai MY, Osanai T (2015) Seawater cultivation of freshwater cyanobacterium *Synechocystis* sp. PCC 6803 drastically alters amino acid composition and glycogen metabolism. Front Microbiol 6: 326

Jablonsky J, Papacek S, Hagemann M (2016) Different strategies of metabolic regulation in cyanobacteria: from transcriptional to biochemical control. Sci Rep 6: 33024

Kaneko T, Sato S, Kotani H, Tanaka A, Asamizu E, Nakamura Y, et al. (1996) Sequence analysis of the genome of the unicellular cyanobacterium *Synechocystis* sp. strain PCC6803. II. Sequence determination of the entire genome and assignment of potential protein-coding regions (supplement). DNA Res 3: 185–209

Kanesaki Y, Suzuki I, Allakhverdiev SI, Mikami K, Murata N (2002) Salt stress and hyperosmotic stress regulate the expression of different sets of genes in *Synechocystis* sp. PCC 6803. Biochem Biophys Res Commun 290: 339–348

Khan RI, Wang Y, Afrin S, Wang B, Liu Y, Zhang X, Chen L, Zhang W, He L, Ma G (2016) Transcriptional regulator PrqR plays a negative role in glucose metabolism and oxidative stress acclimation in *Synechocystis* sp. PCC 6803. Sci Rep 6: 32507

Kizawa A, Kawahara A, Takashima K, Takimura Y, Nishiyama Y, Hihara Y (2017) The LexA transcription factor regulates fatty acid biosynthetic genes in the cyanobacterium *Synechocystis* sp. PCC 6803. Plant J 92: 189–198

Klähn S, Steglich C, Hess WR, Hagemann M (2010a) Glucosylglycerate: a secondary compatible solute common to marine cyanobacteria from nitrogen-poor environments. Environ Microbiol 12: 83–94

Klähn S, Höhne A, Simon E, Hagemann M (2010b) The gene *ssl3076* encodes a protein mediating the salt-induced expression of *ggpS* for the biosynthesis of the compatible solute glucosylglycerol in *Synechocystis* sp. strain PCC 6803. J Bacteriol 192: 4403–4412

Klähn S, Schaal C, Georg J, Baumgartner D, Knippen G, Hagemann M, et al. (2015) The sRNA NsiR4 is involved in nitrogen assimilation control in cyanobacteria by targeting glutamine synthetase inactivating factor IF7. Proc Natl Acad Sci USA 112: E6243–E6252

Kirsch F, Pade N, Klähn S, Hess WR, Hagemann M (2017) The glucosylglycerol degrading enzyme GghA is involved in the acclimation to fluctuating salinities of the cyanobacterium *Synechocystis* sp. strain PCC 6803. Microbiology 163: 1319–1328

Kirsch F, Klähn S, Hagemann M (2019) Salt-regulated accumulation of the compatible solutes sucrose and glucosylglycerol in cyanobacteria and its biotechnological relevance. Front Microbiol 10: 2139

Koksharova O, Schubert M, Shestakov S, Cerff R (1998) Genetic and biochemical evidence for distinct key functions of two highly divergent GAPDH genes in catabolic and anabolic carbon flow of the cyanobacterium *Synechocystis* sp. PCC 6803. Plant Mol Biol 36: 183–194

Kondo K, Geng XX, Katayama M, Ikeuch M (2005). Distinct roles of CpcG1 and CpcG2 in phycobilisome assembly in the cyanobacterium *Synechocystis* sp. PCC 6803. Photosynth Res 84: 269–273

Kopf M, Klähn S, Scholz I, Matthiessen JK, Hess WR, Voß B (2014) Comparative analysis of the primary transcriptome of *Synechocystis* sp. PCC 6803. DNA Res 21: 527–539

Kopf M, Hess WR (2015) Regulatory RNAs in photosynthetic cyanobacteria. FEMS Microbiol Rev 39: 301–315

Kumar L, Futschik ME (2007) Mfuzz: a software package for soft clustering of microarray data. Bioinformation 2: 5–7

Lei H, Chen G, Wang Y, Ding Q, Wei D (2014) Sll0528, a site-2-protease, is critically involved in cold, salt and hyperosmotic stress acclimation of cyanobacterium *Synechocystis* sp PCC 6803. Int J Mol Sci 15: 22678–22693

Levina N, Totemeyer S, Stokes NR, Louis P, Jones MA, Booth IR (1999) Protection of *Escherichia coli* cells against extreme turgor by activation of MscS and MscL mechanosensitive channels: identification of genes required for MscS activity. EMBO J 18: 1730–1737

Li H, Singh AK, McIntyre LM, Sherman LA (2004) Differential gene expression in response to hydrogen peroxide and the putative PerR regulon of *Synechocystis* sp. strain PCC 6803. J Bacteriol 186: 3331–3345

Liu Y, Beyer A, Aebersold R (2016) On the dependency of cellular protein levels on mRNA abundance. Cell 165: 535–550

Liu X, Miao R, Lindberg P, Lindblad P (2019a) Modular engineering for efficient photosyntheteic biosynthesis of 1-butanol from CO_2_ in cyanobacteria. Energy Environ Sci 12: 2765–2777

Liu H, Weis, DA, Zhang MM, Cheng M, Zhang B, Zhang H, Gerstenecker GS, Pakrasi HB, Gross ML, Blankenship RE (2019b) Phycobilisomes harbor FNR L in cyanobacteria. mBio 10: e00669–19

Marin K, Zuther E, Kerstan T, Kunert A, Hagemann M (1998) The *ggpS* gene from *Synechocystis* sp. strain PCC 6803 encoding glucosyl-glycerol-phosphate synthase is involved in osmolyte synthesis. J Bacteriol 180: 4843–4849

Marin K, Huckauf J, Fulda S, Hagemann M (2002) Salt-dependent expression of glucosylglycerol-phosphate synthase, involved in osmolyte synthesis in the cyanobacterium *Synechocystis* sp. strain PCC 6803. J Bacteriol 184: 2870–2877

Marin K, Suzuki I, Yamaguchi K, Ribbeck K, Yamamoto H, Kanesaki Y, Hagemann M, Murata N (2003) Identification of histidine kinases that act as sensors in the perception of salt stress in *Synechocystis* sp. PCC 6803. Proc Natl Acad Sci USA 100: 9061–9066

Marin K, Kanesaki Y, Los DA, Murata N, Suzuki I, Hagemann M (2004) Gene expression profiling reflects physiological processes in salt acclimation of *Synechocystis* sp. strain PCC 6803. Plant Physiol 136: 3290–3300

Matsuhashi A, Tahara H, Ito Y, Uchiyama J, Ogawa S, Ohta H (2015) Slr2019, lipid A transporter homolog, is essential for acidic tolerance in *Synechocystis* sp PCC6803. Photosynth Res 125: 267–277

Mikkat S, Hagemann M (2000) Molecular analysis of the *ggtBCD* operon of *Synechocystis* sp. strain PCC 6803 encoding the substrate-binding protein and the transmembrane proteins of an ABC transporter. Arch Microbiol 174: 273–282

Mikkat S, Milkowski C, Hagemann M (2000) The gene *sll*0273 of the cyanobacterium *Synechocystis* sp. strain PCC 6803 encodes a protein essential for growth at low Na^+^/K^+^ ratios. Plant Cell Environm 23: 549–559

Mitschke J, Georg J, Scholz I, Sharma CM, Dienst D, Bantscheff J, et al. (2011) An experimentally anchored map of transcriptional start sites in the model cyanobacterium *Synechocystis* sp. PCC6803. Proc Natl Acad Sci USA 108: 2124–2129

Novak JF, Stirnberg M, Roenneke B, Marin K (2011) A novel mechanism of osmosensing, a salt-dependent protein-nucleic acid interaction in the cyanobacterium *Synechocystis* Species PCC 6803. J Biol Chem 286: 3235–3241

Nowaczyk MM, Krause K, Mieseler M, Sczibilanski A, Ikeuchi M, Rögner M (2012) Deletion of *psbJ* leads to accumulation of Psb27-Psb28 photosystem II complexes in *Thermosynechococcus elongatus*. Biochim Biophys Acta 1817: 1339–1345

Pade N, Compaoré J, Klähn S, Stal LJ, Hagemann M (2012) The marine cyanobacterium *Crocosphaera watsonii* WH8501 synthesizes the compatible solute trehalose by a laterally acquired OtsAB fusion protein. Environ Microbiol 14: 1261–1271

Pade N, Hagemann M (2014) Salt acclimation of cyanobacteria and their application in biotechnology. Life 5: 25–49

Pade N, Michalik D, Ruth W, Belkin N, Hess WR, Berman-Frank I, Hagemann M (2016) Trimethylated homoserine functions as the major compatible solute in the globally significant oceanic cyanobacterium *Trichodesmium*. Proc Natl Acad Sci USA 113: 13191–13196

Pade N, Mikkat S, Hagemann M (2017) Ethanol, glycogen and glucosylglycerol represent competing carbon pools in ethanol-producing cells of *Synechocystis* sp. PCC 6803 under high-salt conditions. Microbiology 163: 300–307

Pappesch R, Warnke P, Mikkat S, Normann J, Wisniewska-Kucper A, Huschka F, Wittmann M, Khani A, Schwengers O, Oehmcke-Hecht S, Hain T, Kreikemeyer B, Patenge N. (2017) The regulatory small RNA *marS* supports virulence of *Streptococcus pyogenes*. Sci Rep 7: 12241

Pattanayak GK, Liao Y, Wallace EWJ, Budnik B, Drummond DA, Rust MJ (2020) Daily cycles of reversible protein condensation in cyanobacteria. Cell Rep 32: 108032

Pereira SB, Santos M, Leite JP, Flores C, Eisfeld C, Büttel Z, Mota R, Rossi F, De Philippis R, Gales L, Morais-Cabral JH, Tamagnini P (2019) The role of the tyrosine kinase Wzc (Sll0923) and the phosphatase Wzb (Slr0328) in the production of extracellular polymeric substances (EPS) by *Synechocystis* PCC 6803. Microbiol open 8: e00753

Perozo E, Kloda A, Cortes DM, Martinac B. (2001) Site-directed spin-labeling analysis of reconstituted MscL in the closed state. J Gen Physiol 118: 193–206

Pinto FL, Thapper A, Sontheim W, Lindblad P (2009) Analysis of current and alternative phenol based RNA extraction methodologies for cyanobacteria. BMC Mol Biol 10: 79

Prakash JS, Sinetova M, Zorina A, Kupriyanova E, Suzuki I, Murata N, Los DA (2009) DNA supercoiling regulates the stress-inducible expression of genes in the cyanobacterium *Synechocystis*. Mol Biosyst 5: 1904–1912

Qiao J, Huang S, Te R, Wang J, Chen L, Zhang W (2013) Integrated proteomic and transcriptomic analysis reveals novel genes and regulatory mechanisms involved in salt stress responses in *Synechocystis* sp. PCC 6803. Appl Microbiol Biotechnol 97: 8253–8264

Qiao Y, Wang W, Lu X (2020) Engineering cyanobacteria as cell factories for direct trehalose production from CO_2_. Metab Eng 62: 161–171

Reed RH, Borowitzka LJ, Mackay MA, Chudek JA, FosterR, Warr SRC, et al. (1986) Organic solute accumulation in osmotically stressed cyanobacteria. FEMS Microbiol Lett 39: 51–56

Reed RH, Stewart WDP (1985) Osmotic adjustment and organic solute accumulation in unicellular cyanobacteria from freshwater and marine habitats. Mar Biol 88: 1–9

Riediger M, Kadowaki T, Nagayama R, Georg J, Hihara Y, Hess WR (2019) Biocomputational analyses and experimental validation identify the regulon controlled by the redox-responsive transcription factor RpaB. iScience 15: 316–331

Riediger M, Spät P, Bilger R, Voigt K, Macek B, Hess WR (2021) Analysis of a photosynthetic cyanobacterium rich in internal membrane systems via gradient profiling by sequencing (Grad-seq). Plant Cell 33: 248–269

Ritter SPA, Lewis AC, Vincent SL, Lo LL, Cunha APA, Chamot D, Ensminger I, Espie GS, Owttrim GW (2020) Evidence for convergent sensing of multiple abiotic stresses in cyanobacteria. Biochim Biophys Acta Gen Subj 1864: 129462

Rübsam H, Kirsch F, Reimann V, Erban A, Kopka J, Hagemann M, Hess WR, Klähn S (2018) The iron-stress activated RNA 1 (IsaR1) coordinates osmotic acclimation and iron starvation responses in the cyanobacterium *Synechocystis* sp. PCC 6803. Environ Microbiol 20: 2757–2768

Sakurai I, Stazic D, Eisenhut M, Vuorio E, Steglich C, Hess WR, Aro EM (2012) Positive regulation of *psbA* gene expression by cis-encoded antisense RNAs in *Synechocystis* sp. PCC 6803. Plant Physiol 160: 1000–1010

Scholz I, Lange SJ, Hein S, Hess WR, Backofen R (2013) CRISPR-Cas systems in the cyanobacterium *Synechocystis* sp. PCC6803 exhibit distinct processing pathways involving at least two Cas6 and a Cmr2 protein. PLoS One 8: e56470

Schwarz D, Orf I, Kopka J, Hagemann M (2013) Recent applications of metabolomics toward cyanobacteria. Metabolites 3: 72–100

Shcolnick S, Shaked Y, Keren N (2007) A role for mrgA, a DPS family protein, in the internal transport of Fe in the cyanobacterium *Synechocystis* sp. PCC6803. Biochim Biophys Acta 1767: 814–819

Shapiguzov A, Lyukevich AA, Allakhverdiev SI, Sergeyenko TV, Suzuki I, Murata N, Los DA (2005) Osmotic shrinkage of cells of *Synechocystis* sp. PCC 6803 by water efflux via aquaporins regulates osmostress-inducible gene expression. Microbiology 151: 447–455

Shi L, Bischoff KM, Kennelly PJ (1999) The *icfG* gene cluster of *Synechocystis* sp. strain PCC 6803 encodes an Rsb/Spo-like protein kinase, protein phosphatase, and two phosphoproteins. J Bacteriol 181: 4761–4767

Shoumskaya MA, Paithoonrangsarid K, Kanesaki Y, Los DA, Zinchenko VV, Tanticharoen M, Suzuki I, Murata N (2005) Identical Hik-Rre systems are involved in perception and transduction of salt signals and hyperosmotic signals but regulate the expression of individual genes to different extents in *Synechocystis*. J Biol Chem 280: 21531–21538

Silva JC, Gorenstein MV, Li GZ, Vissers JPC, Geromanos SJ (2006) Absolute quantification of proteins by LCMSE: A virtue of parallel MS acquisition. Mol Cell Proteomics 5: 144–156

Spät P, Barske T, Maček B, Hagemann M (2021) Alterations in the CO_2_ availability induce alterations in the phospho-proteome of the cyanobacterium *Synechocystis* sp. PCC 6803. New Phytol 231: 1123–1137

Stanier RY, Kunisawa R, Mandel M, Cohen-Bazire G (1971) Purification and properties of unicellular blue-green algae (order *Chroococcales*). Bacteriol Rev 35: 171–205

Stokes NR, Murray HD, Subramaniam C, Gourse RL, Louis P, Bartlett W, Miller S, Booth IR (2003) A role for mechanosensitive channels in survival of stationary phase: regulation of channel expression by RpoS. Proc Natl Acad Sci USA 100: 15959–15964

Summerfield TC, Sherman LA (2008) Global transcriptional response of the alkali-tolerant cyanobacterium *Synechocystis* sp. strain PCC 6803 to a pH 10 environment. Appl Environ Microbiol 74: 5276–5284

Takashima K, Nagao S, Kizawa A, Suzuki T, Dohmae N, Hihara Y (2020) The role of transcriptional repressor activity of LexA in salt-stress responses of the cyanobacterium *Synechocystis* sp. PCC 6803. Sci Rep 10: 17393

Toyoshima M, Tokumaru Y, Matsuda F, Shimizu H (2020) Assessment of protein content and phosphorylation level in *Synechocystis* sp. PCC 6803 under various growth conditions using quantitative phosphoproteomic analysis. Molecules 25: E3582

Tyystjärvi T, Huokko T, Rantamäki S, Tyystjärvi E (2013) Impact of different group 2 sigma factors on light use efficiency and high salt stress in the cyanobacterium *Synechocystis* sp. PCC 6803. PLoS One 8: e63020

Uchiyama J, Asakura R, Moriyama A, Kubo Y, Shibata Y, Yoshino Y, Tahara H, Matsuhashi A, Sato S, Nakamura Y, Tabata S, Ohta H (2014) Sll0939 is induced by Slr0967 in the cyanobacterium *Synechocystis* sp. PCC6803 and is essential for growth under various stress conditions. Plant Physiol Biochem 81: 36–43

Vinnemeier J, Hagemann M (1999) Identification of salt-regulated genes in the genome of the cyanobacterium Synechocystis sp. strain PCC 6803 by subtractive RNA hybridization. Arch Microbiol 172: 377–386

Vizcaíno JA, Cote’ RG, Csordas A, Dianes JA, Fabregat A, Foster JM, Griss J, Alpi E, Birim M, Contell J, et al. (2013) The Proteomics Identifications (PRIDE) database and associated tools: status in 2013. Nucl Acid Res 41: D1063–D1069

Wang HL, Postier BL, Burnap RL (2002) Polymerase chain reaction-based mutageneses identify key transporters belonging to multigene families involved in Na^+^ and pH homeostasis of *Synechocystis* sp. PCC 6803. Mol Microbiol 44: 1493–1506

Wegener KM, Singh AK, Jacobs JM, Elvitigala T, Welsh EA, Keren N, Gritsenko MA, Ghosh BK, Camp DG 2nd, Smith RD, et al. (2010) Global proteomics reveal an atypical strategy for carbon/nitrogen assimilation by a cyanobacterium under diverse environmental perturbations. Mol Cell Proteomics 9: 2678–2689

Whitton BA, Potts M (2000) The ecology of cyanobacteria. Their diversity in time and space. Kluwer Academic Publishers, Dordrecht, The Netherlands

Wu W, Du W, Gallego RP, Hellingwerf KJ, van der Woude AD, Branco Dos Santos F (2020) Using osmotic stress to stabilize mannitol production in *Synechocystis* sp. PCC6803. Biotechnol Biofuels 13: 117

Zhan J, Steglich C, Scholz I, Hess WR, Kirilovsky D (2021) Inverse regulation of light harvesting and photoprotection mediated by a 3’end-derived sRNA in cyanobacteria. Plant Cell 33: 358–380

Zuther E, Schubert H, Hagemann M. 1998. Mutation of a gene encoding a putative glycoprotease leads to reduced salt tolerance, altered pigmentation, and cyanophycin accumulation in the cyanobacterium *Synechocystis* sp. strain PCC 6803. J Bacteriol 180: 1715–1722

