## Supplementary material for "Integrative analysis of the salt stress response in cyanobacteria": Suppl. Methods and Figures

### Detailed description of proteomic methods

#### Subcellular fractionation

Cells from four biological replicates of salt-acclimated and control cultures, respectively, were broken with glass beads using Precellys 24 homogenizer (peqLab Biotechnologie GmbH, Erlangen, Germany) in non-denaturing buffer containing 10 mM Tris/HCl, pH 7.4, 138 mM NaCl, 2.7 mM KCl, 1 mM $MgCl_2$. An aliquot of each cell extract was saved as total protein immediately after cell disruption. The cell debris was separated from the remaining cell extracts by low speed centrifugation at 750 g (5 min, 4 °C). Debris from the biological replicates was pooled, washed once with non-denaturing buffer and stored as debris fraction. After removing debris, the cell extracts were centrifuged at 22,000 g, 4°C for 90 min to obtain the soluble and membrane-enriched fractions. The membrane-enriched fractions were 2-times washed with high salt (1 M KCl) and high pH (0.1 M $Na_2CO_3$) buffers to deplete loosely bound proteins. Protein concentration was measured using the Bio-Rad protein assay (Bio-Rad Munich, Germany).

**In-solution digestion of proteins**

Aliquots of the protein fractions were supplemented with stock solutions of ammonium bicarbonate (ABC), sodium deoxycholate (SDC) and dithiothreitol (DTT) to obtain samples containing between 40 and 75 µg protein in 100 µl extraction buffer (50 mM ABC, 1.5% SDC, 10 mM DTT). The protein samples were incubated at 95 °C for 5 min and subsequently sonicated for 10 min using a bath sonicator. Alkylation was performed with 15 mM iodoacetamide for 20 min at room temperature. Thereafter, 50 mM ABC and sequencing grade trypsin (Promega) were added to obtain an enzyme/protein ratio of 1:50 in a final volume of 320 µl. Digestion was performed at 37°C for about 16 h. SDC was removed from the digest solutions using the phase transfer surfactant method (Matsuda et al. 2008) according to Pappesch et al. (2017) by adding 320 µl of ethylacetate and 6.5 µl of 25% trifluoroacetic acid, rigorous mixing for 3 min and subsequent centrifugation at 12,000 g for 10 min to obtain aqueous and organic phases. 200 µl of the aqueous phase was collected using a gel loading tip. Finally, the peptide solutions were desalted with OASIS HLB 1cc Vac Cartridges (Waters, Manchester, UK), concentrated using a centrifugal evaporator and dissolved with 2% acetonitrile in 0.1% formic acid for mass spectrometric analysis. Peptide concentrations were measured using the Invitrogen Qubit protein assay kit (Thermo Fisher Scientific).

**Analysis by nanoLC-HDMS$^E$**

LC-HDMS$^E$ analyses were carried out using a nanoAcquity UPLC system (Waters) coupled to a Waters Synapt G2-S mass spectrometer via a NanoLockSpray ion source as described (Pade et al., 2017). Mobile phase A contained 0.1% formic acid in water and mobile phase B contained 0.1% formic acid in acetonitrile. Peptide samples corresponding to approximately 150 ng (soluble fraction), 180 ng (total extract) and 360 ng (membrane and debris fraction) of digested protein, supplemented with 40 fmol of Hi3 Phos B standard for protein absolute quantification (Waters), were trapped and desalted using a precolumn (ACQUITY UPLC Symmetry C18, 5 µm, 180 µm x 20 mm, Waters) at a flow rate of 10 µl/min for 4 min with 99.9% A. Peptides were separated on an analytical column (ACQUITY UPLC HSS T3, 1.8 µm, 75 µm x 250 mm, Waters) at a flow rate of 300 nl/min using a gradient from 3% to 32% B over 150 min. The column temperature was maintained at 35°C. The Synapt G2-S instrument was operated in data-independent mode with ion-mobility separation as an additional dimension of separation (referred to as HDMS$^E$). By executing alternate scans at low and elevated collision energy (CE) of each 0.6 sec, information on precursor and fragment ions, respectively, was acquired. In low-energy MS mode acquisitions were performed at constant CE of 4 eV whereas drift time-dependent CE settings (Distler et al., 2014) were applied in elevated-energy MS mode. As a reference compound, 100 fmol/µl [Glu1]-fibrinopeptide B was delivered at 500 nl/min to the reference sprayer of the NanoLockSpray source. Lock spray was acquired once every 30 s for a 1 s period. Single measurements of the four biological replicates of the total extract, the soluble fraction and the membrane-enriched fraction were carried out, while the pooled samples of the debris fraction were measured in triplicate.

**NanoLC-HDMS$^E$ data processing, protein identification and quantification**

Progenesis QI for Proteomics version 4.1 (Nonlinear Dynamics, Newcastle upon Tyne, UK) was used for raw data processing, protein identification and label free quantification. The raw data from the measurements of the total extracts and each of the three protein fractions were analyzed separately. Normalization and matching of features (accurate mass retention time tags) between runs was performed to compensate for between-run variation in the LC separation and to avoid missing values in quantification. For the protein identification a database containing 3507 protein sequences from *Synechocystis* 6803 (UniProt release 2016_07) appended with the sequences of rabbit phosphorylase B (P00489) and porcine trypsin was compiled. Two missing cleavage sites were allowed, oxidation of methionine residues was considered as variable modification, and carbamidomethylation of cysteine's as fixed modification. The false discovery rate was set to 1%. Peptides were required to be identified by at least three fragment ions and proteins by at least two peptides and six fragment ions. Subsequently peptide ion data were filtered to retain only peptide ions that met the following criteria: (i) identified at least in two samples within the dataset, (ii) minimum ion score of 5.7 (total extract and the soluble fraction) or 5.9 (membrane-enriched and debris fraction), (iii) mass error below 12.0 ppm, and (iiii) at least 6 amino acid residues in length. Identifications based on charge state deconvolution were removed.

Proteins were quantified by the absolute quantification Hi3 method using Hi3 Phos B Standard (Waters) as reference (Silva et al., 2006). Results were given as fmol on column. The peptide expression profiles of selected proteins were reviewed manually in order to remove outlier peptides whose expression differed significantly from the other peptides of the same protein. Only proteins identified in one fraction by at least two unique peptides were included in the quantitative analysis. Protein abundance changes between salt-acclimated and control cells by a factor of at least 1.5, accompanied by ANOVA *p*-values < 0.05 were regarded as significant. To determine a combined fold change value for the total extract and the three subcellular fractions, first a weighted fold change value for the three subcellular fractions was calculated by summing up the corresponding protein amounts (measured as fmol on column) and dividing the value from salt-acclimated cells by the value from controls. Since the debris fraction contained only a minor part of about 10% of the total protein, the protein amounts of the debris fraction were divided by ten. The calculation of this weighted fold change is shown in the following formula, where the abbreviation MA denotes the average protein abundance from the replicate samples: Weighted fold change$_{fractions}$ = (MA 4%$_{membrane}$ + MA 4%$_{soluble}$ +1/10 MA 4%$_{debris}$)/(MA 0%$_{membrane}$ + MA 0%$_{soluble}$ +1/10 MA 0%$_{debris}$). In a second step, the final combined fold change value was calculated as the average of the weighted fold change from the subcellular fractions and the fold change of the total extract. If a fold change for one of the individual fractions had a value > 1.5, but was accompanied by a $p$-value > 0.05, these data were excluded from the calculations.

### Supplementary Figures

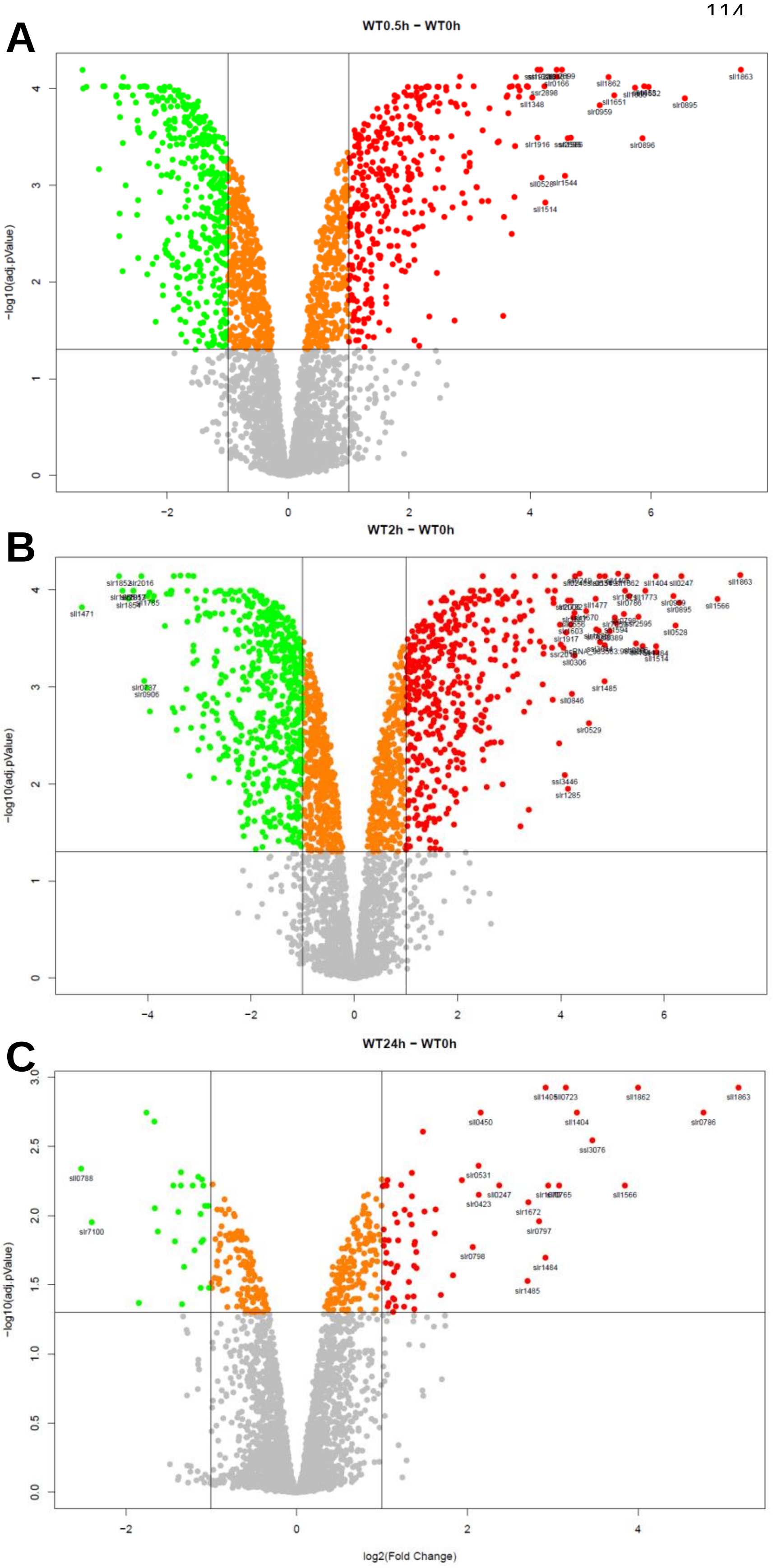


**Supplementary Figure S1. Overall changes in the expression of protein-coding genes between control and salt-stressed cells.** Volcano plots show the -$\log_{10}$ Anova p-value against the $\log_2$ fold change (salt-acclimated/control) for mRNAs in *Synechocystis* 6803 cells grown in NaCl-free BG11 medium (0 h) and medium supplemented with 4% NaCl for 0.5 h (A), 2 h (B) or 24 h (C). Each mRNA is represented as a dot. Transcripts with statistically significant differential expression ($\log_2$ fold change ≥│1│, *P* value ≤ 0.05) are located in the top right and top left quadrants.

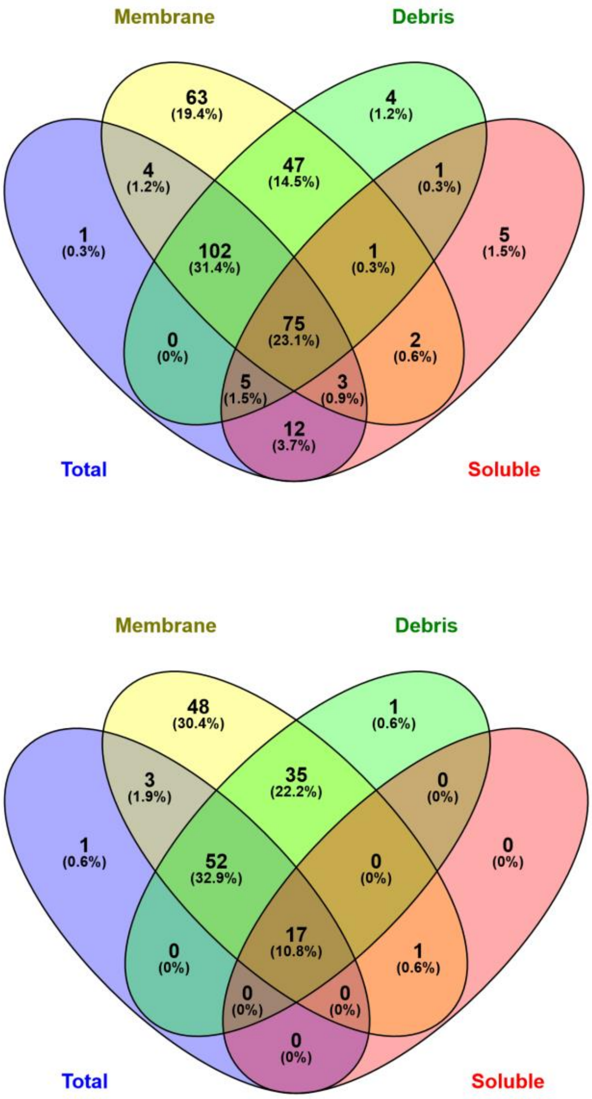


**Figure S2. Venn diagrams showing numbers and percentage of identified proteins** with at least one (A) or two (B) transmembrane helices in the total protein extract and in the subcellular fractions.

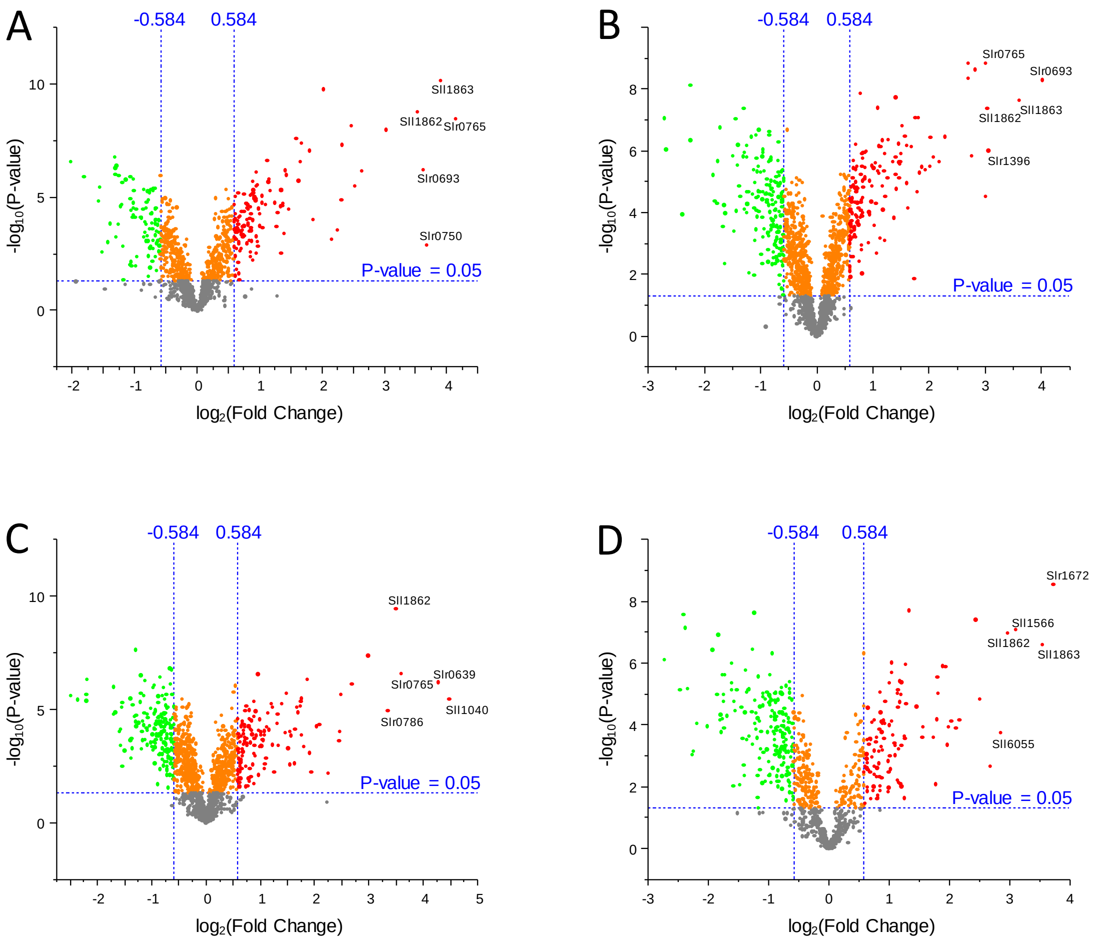


**Figure S3. Overall changes in the proteome between control and salt-acclimated cells.** Volcano plots are shown that indicate differential abundances of proteins. The $-\log_{10}$ Anova p-value is plotted against the $\log_2$ fold change (salt-acclimated/control) for proteins identified in the total protein extract (A), the membrane-enriched fraction (B), the debris fraction (C), and the soluble fraction (D) of protein extracts from *Synechocystis* 6803 cells grown in NaCl-free BG11 medium (control) or medium supplemented with 4% NaCl for 7 days. Each protein is represented as a dot. Proteins with statistically significant differential expression (absolute fold change of >1.5 or <-1.5*, p-value <0.05) are located in the top right and top left quadrants. *Please note that the x-axis is $\log_2$ scaled and the fold change threshold is given at a value of 0.584 accordingly.

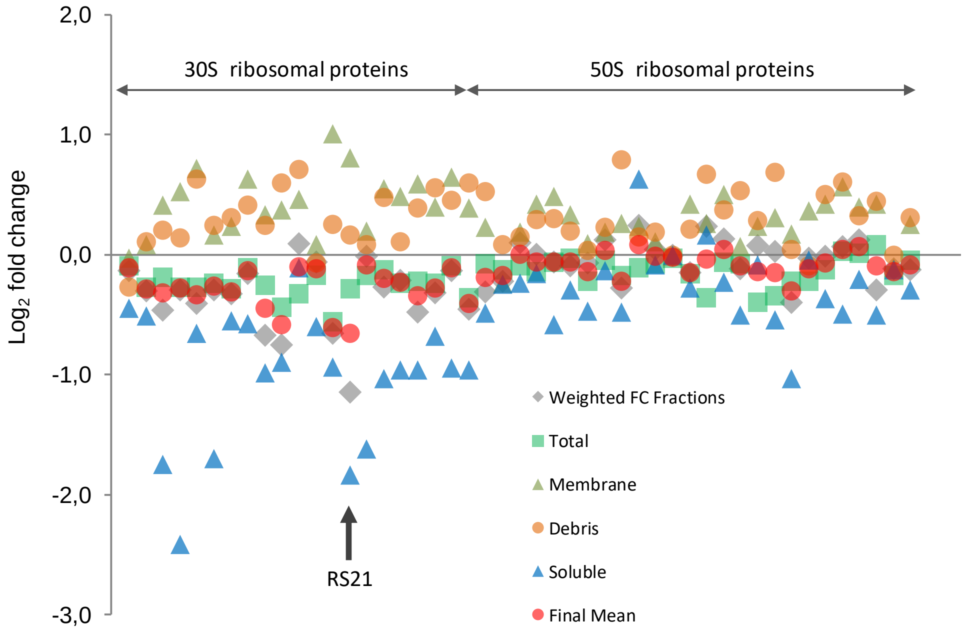


**Figure S4. Growth condition-dependent distribution of ribosomal proteins to the subcellular fractions.** For each ribosomal protein, $log_2$ fold change values (salt-acclimated/control) from the three subcellular fractions, the weighted fold change calculated from the individual fractions, the fold change from the total protein extract, and the finally used combined fold changes are plotted vertically aligned. The combined fold change (final mean) represents the average of the weighted fold change from fractions and the fold change from the total protein extract. Despite large differences between the fold change values from the subcellular fractions, the weighted fold change and the fold change from the total protein extract are close together with the exception of the RS21 protein indicated by an arrow. This protein and proteins showing similar behaviour were excluded from further quantitative analysis.

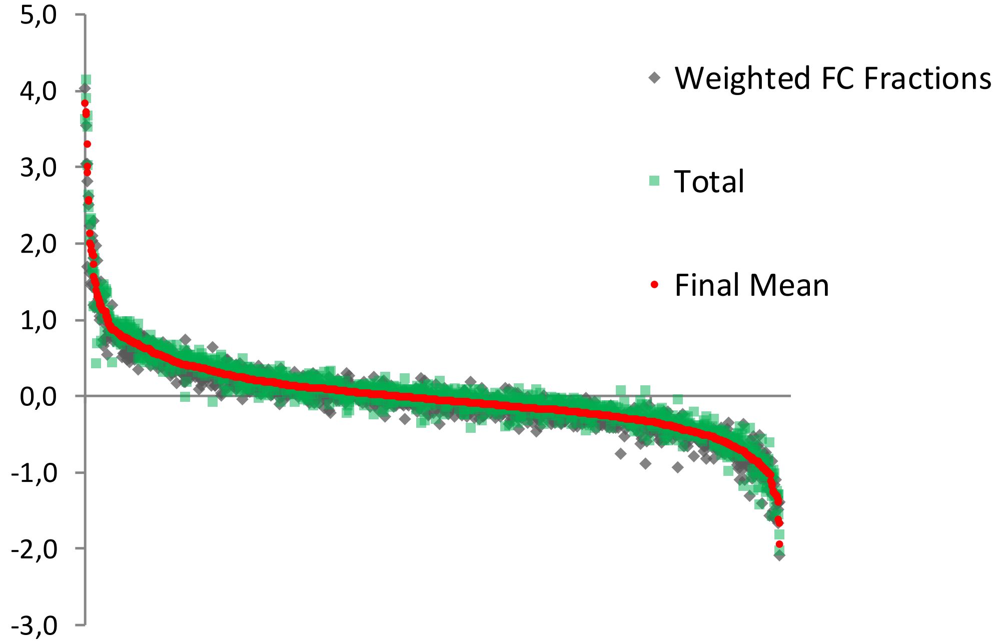


**Figure S5.** Correlation between the weighted fold change value calculated from the subcellular fractions and the fold change value from the total protein extract. The finally used combined fold change value (final mean) was calculated as the average of the weighted fold change from the subcellular fractions and the fold change of the total protein extract.

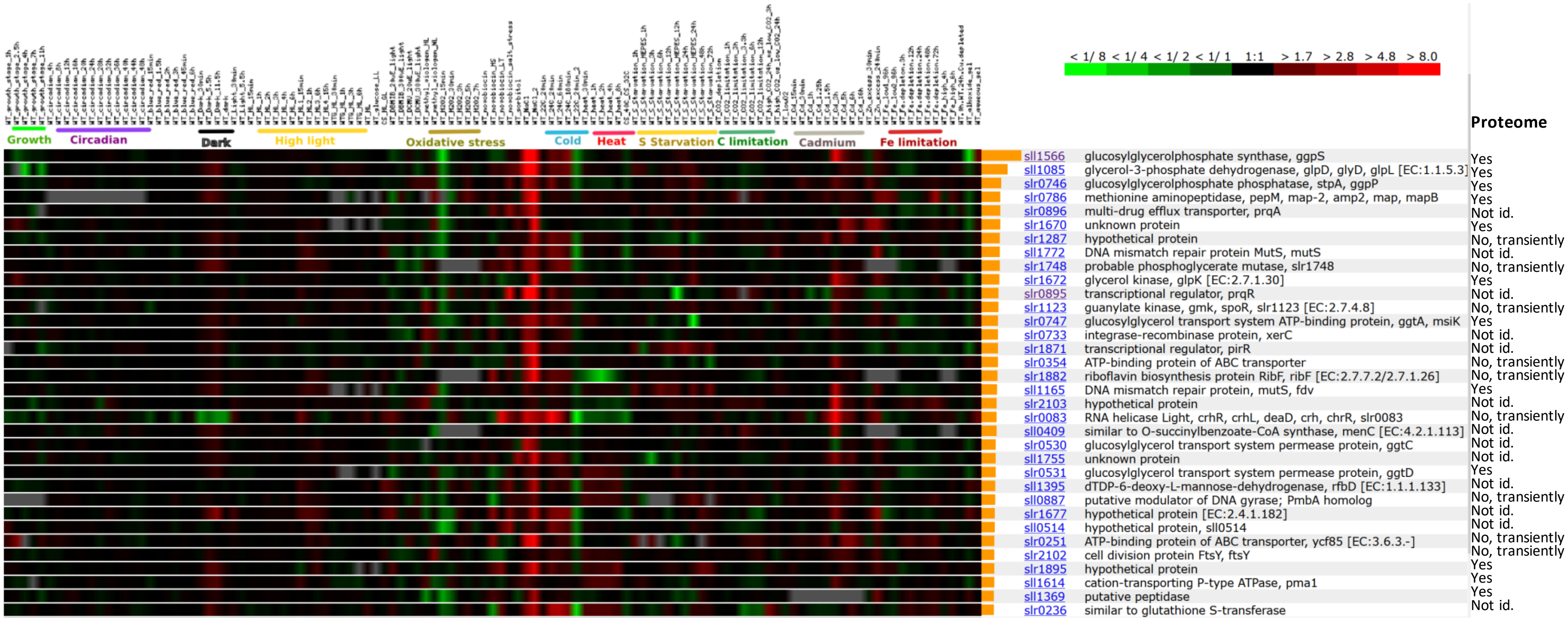


**Figure S6**: **Co-expression of salt-specific mRNAs under different environmental conditions using the data base CyanoExpress (Hernandez-Prieto and Futschik, 2012).** CyanoEXpress: A web database for exploration and visualization of the integrated transcriptome of cyanobacterium Synechocystis sp. PCC6803. Bioinformation 8:634-638). The top 34 genes are displayed that show similar stress-regulated patterns as the salt-regulated gene *ggpS* (*sll1566*). The corresponding changes at proteome level are summarized in the right column (Yes – protein also salt-induced; No – protein not stably salt-induced; Not id. – not identified in the proteome data set; transiently, gene expression is only transiently stimulated after salt shock).

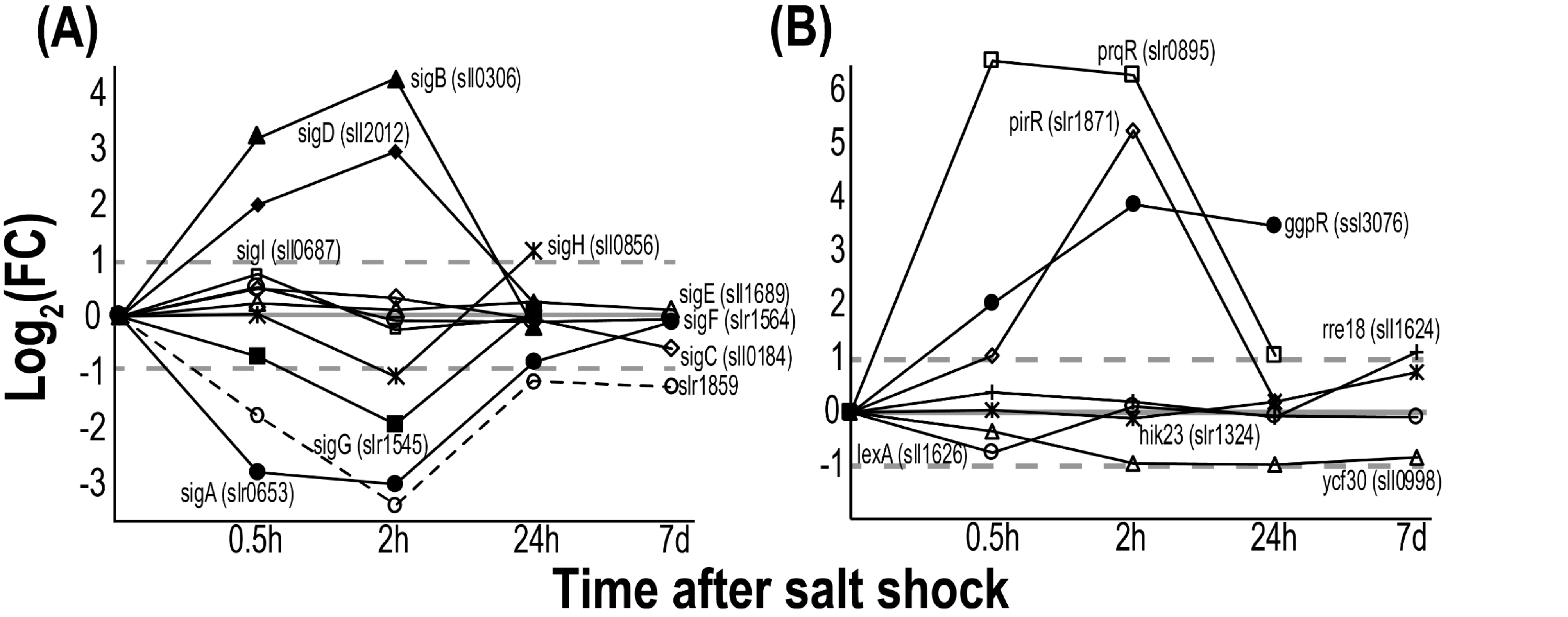


**Figure S7**: **Expression of regulatory proteins in *Synechocystis* 6803 at different time points after addition of 4% NaCl.** The data at time points 0.5 – 24 h represent mRNA values from the microarray data set, while the 7 day expression levels shows the alteration of protein abundances (when identified). **A.** Annotated sigma factor genes. **B.** Genes for selected transcriptional factors.

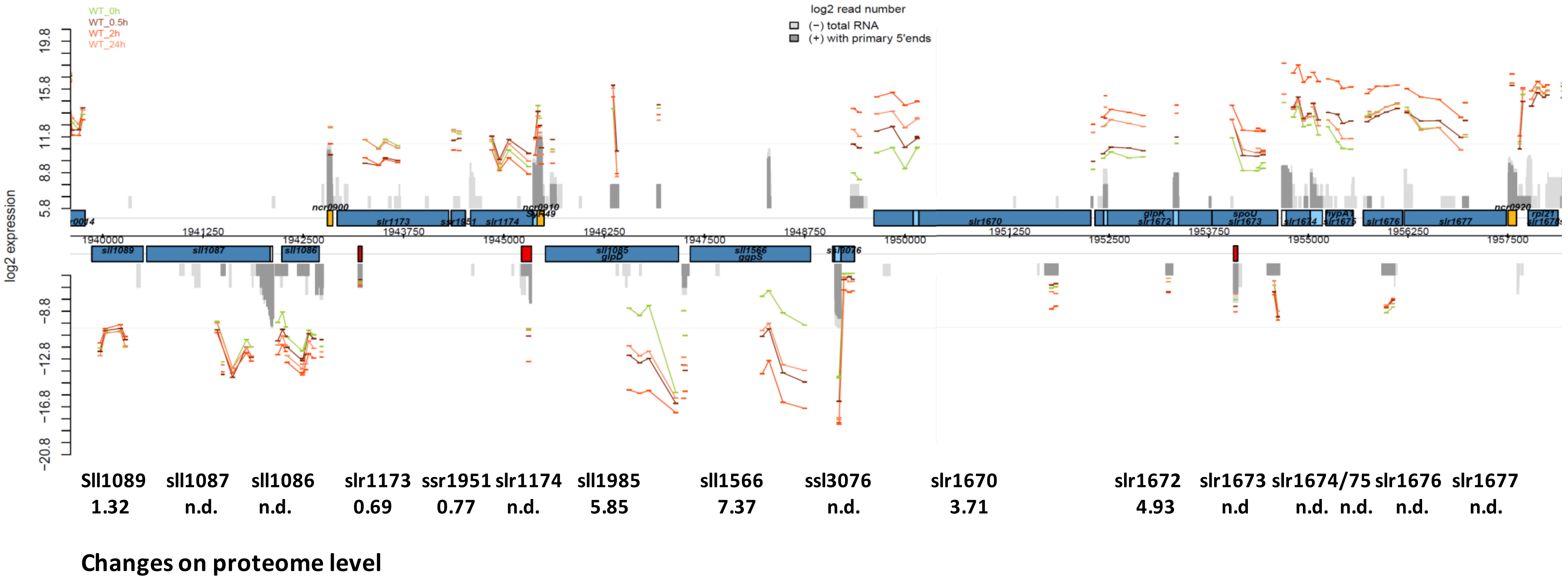


**Figure S8**: **Salt-regulated chromosomal region comprising coordinately up-regulated genes such as *ggpS* (*sll1566*).** The upper panel shows the genome plot and added microarray data with $log_2$ changed expression (low salt-grown control in green, different salt sampling points in reddish color). Grey regions show dRNAseq results (Kirsch et al., 2017), i.e. dark grey columns display active TSSs. The lower panel displays the corresponding proteome changes given as fold changes.

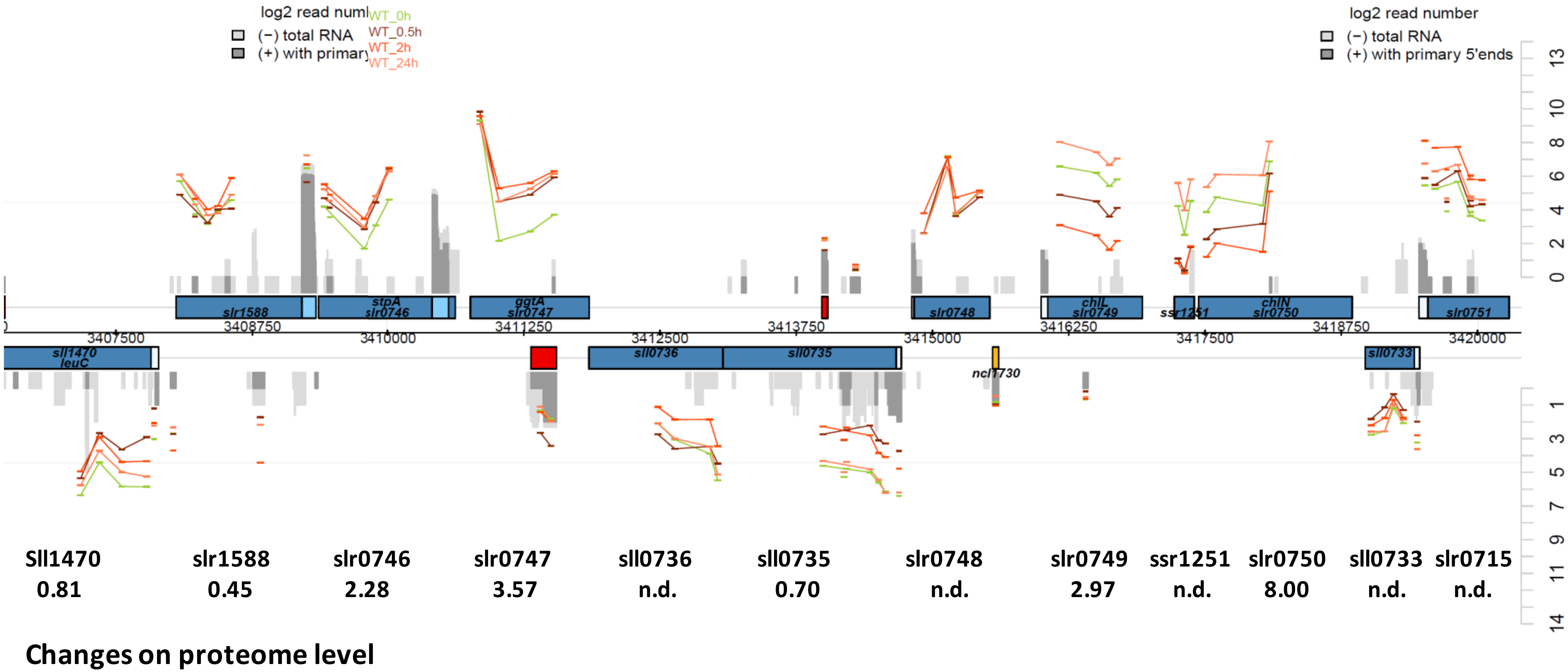


**Figure S9**: **Salt-regulated chromosomal region comprising coordinately up-regulated genes such as *ggpP* (*stpA*, *slr0746*).** The upper panel shows the genome plot and added microarray data with $log_2$ changed expression (low salt-grown control in green, different salt sampling points in reddish color). Grey regions show dRNAseq results (Kirsch et al., 2017), i.e. dark grey columns display active TSSs. The lower panel displays the corresponding proteome changes given as fold changes.

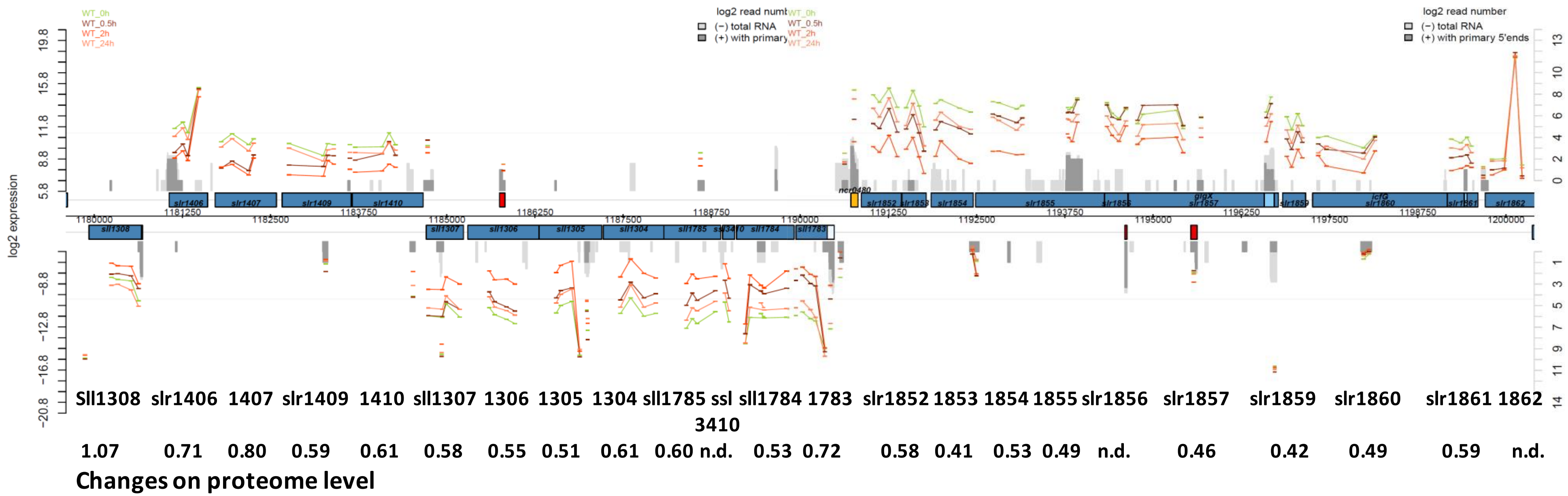


**Figure S10**: **Salt-regulated chromosomal region comprising coordinately down-regulated genes such as *icfG* (*slr1860*).** The upper panel shows the genome plot and added microarray data with $log_2$ changed expression (low salt-grown control in green, different salt sampling points in reddish color). Grey regions show dRNAseq results (Kirsch et al., 2017), i.e. dark grey columns display active TSSs. The lower panel displays the corresponding proteome changes given as fold changes.

### Additional references

Distler U, Kuharev J, Navarro P, et al., (2014) Drift time-specific collision energies enable deep-coverage data-independent acquisition proteomics. Nature Methods 11, 167-170.

Kirsch F, Pade N, Klähn S, Hess WR, Hagemann M (2017) The glucosylglycerol degrading enzyme GghA is involved in the acclimation to fluctuating salinities of the cyanobacterium *Synechocystis* sp. strain PCC 6803. Microbiology 163: 1319-1328.

Masuda T, Tomita M, Ishihama Y (2008). Phase transfer surfactant-aided trypsin digestion for membrane proteome analysis. J Proteome Res 7, 731–740.
